# Using Deep Clustering to Improve fMRI Dynamic Functional Connectivity Analysis

**DOI:** 10.1101/2021.12.14.472680

**Authors:** Arthur P C Spencer, Marc Goodfellow

## Abstract

Dynamic functional connectivity (dFC) analysis of resting-state fMRI data is commonly performed by calculating sliding-window correlations (SWC), followed by k-means clustering in order to assign each window to a given state. Studies using synthetic data have shown that k-means performance is highly dependent on sliding window parameters and signal-to-noise ratio. Additionally, sources of heterogeneity between subjects may affect the accuracy of group-level clustering, thus affecting measurements of dFC state temporal properties such as dwell time and fractional occupancy. This may result in spurious conclusions regarding differences between groups (e.g. when comparing a clinical population to healthy controls). Therefore, is it important to quantify the ability of k-means to estimate dFC state temporal properties when applied to cohorts of multiple subjects, and to explore ways in which clustering performance can be maximised.

Here, we explore the use of dimensionality reduction methods prior to clustering in order to map high-dimensional data to a lower dimensional space, providing salient features to the subsequent clustering step. We assess the use of deep autoencoders for feature selection prior to applying k-means clustering to the encoded data. We compare this deep clustering method to feature selection using principle component analysis (PCA), uniform manifold approximation and projection (UMAP), as well as applying k-means to the original feature space using either L1 or L2 distance. We provide extensive quantitative evaluation of clustering performance using synthetic datasets, representing data from multiple heterogeneous subjects. In synthetic data we find that deep clustering gives the best performance, while other approaches are often insufficient to capture temporal properties of dFC states. We then demonstrate the application of each method to real-world data from human subjects and show that the choice of feature selection method has a significant effect on group-level measurements of state temporal properties. We therefore advocate for the use of deep clustering as a precursor to clustering in dFC.

## 1. Introduction

Functional connectivity (FC) analysis is used to characterise and quantify the spatiotemporal patterns of brain activity, measured non-invasively using functional magnetic resonance imaging (fMRI). In these analyses, statistical similarities (e.g. correlations) in the blood oxygen level dependent (BOLD) signal between pairs of brain regions, or “nodes”, are used as edges in the construction of FC networks. This facilitates the use of graph theory to quantify whole-brain dynamics (Bassett and Sporns, 2017; Bullmore and Sporns, 2009; Bullmore and Bassett, 2011; Medaglia et al., 2015). More recently, dynamic functional connectivity (dFC) has been widely adopted to investigate the time-varying organisation of resting-state brain activity (Calhoun et al., 2014; Cohen, 2018; Hutchison et al., 2013; Karahanoğlu and Van De Ville, 2017; Lurie et al., 2020; Preti et al., 2017). A common approach is to calculate sliding-window correlations (SWC), resulting in a set of FC matrices that can then be clustered into sets of repetitively occurring FC patterns, or “states” (Allen et al., 2014; Calhoun et al., 2014), as summarised in Fig. 1. For a review, see Preti et al. (2017). The spatiotemporal dynamics of the brain are then quantified in terms of the stability and variability of each dFC state, using statistics such as dwell time (the average duration the state is occupied before a state change) and fractional occupancy (the proportion of time spent in a given state).

**Figure 1:**
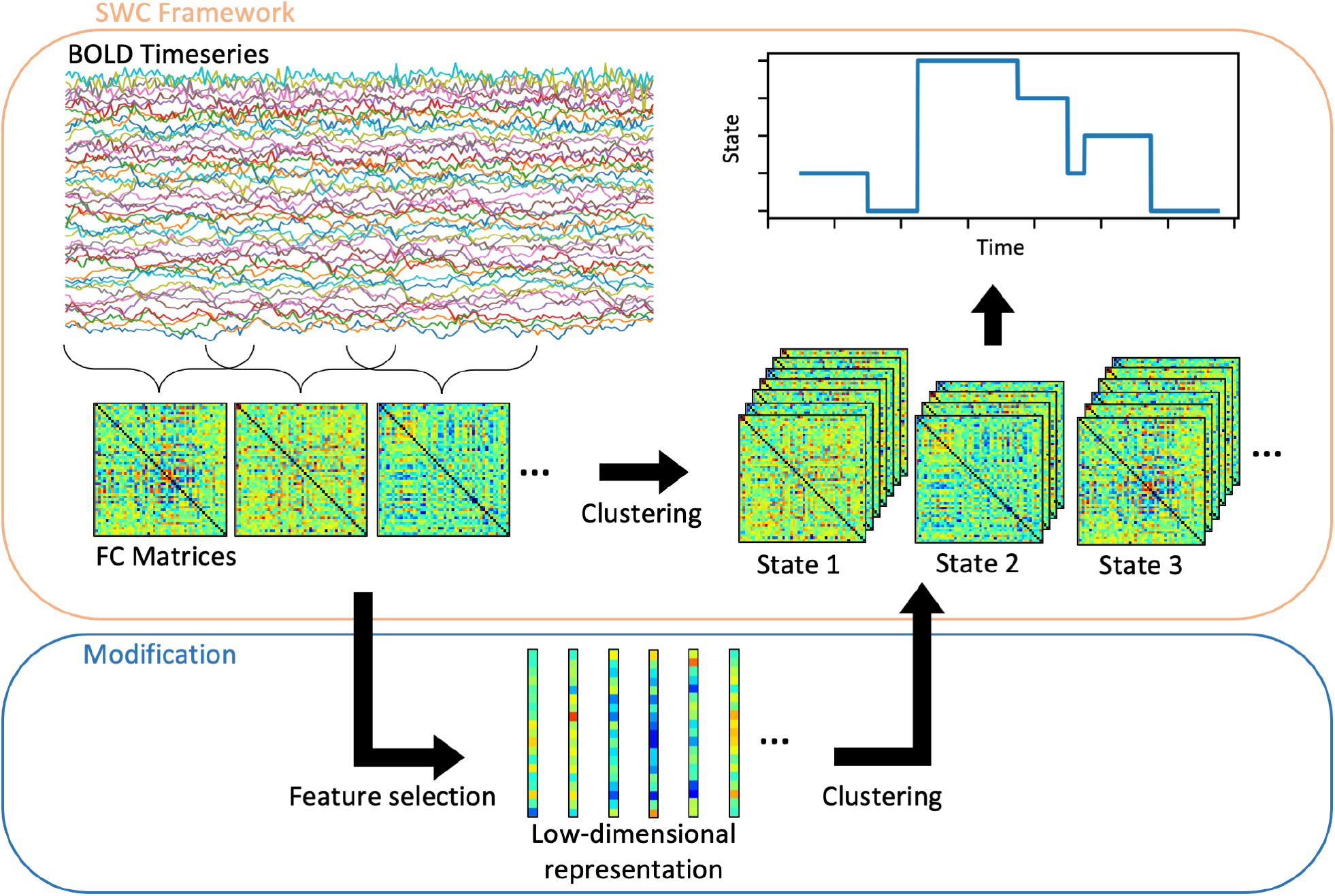
Overview of the SWC framework, including constructing FC matrices from BOLD fMRI data, followed by clustering in order to determine dFC states. Also shown is the modification to this framework assessed in this study, which consists of applying feature selection to the FC matrices before clustering the low-dimensional representation of the data.

Thus, an important assumption underlying dFC analysis is the ability to segment the dynamics of the brain into states with particular spatial correlation patterns, and that these patterns remain stationary for some time period shorter than the length of the scan (analogous to EEG microstates (Michel and Koenig, 2018)). This is typically achieved in practice using clustering methods (e.g. k-means) to identify which of the windowed FC matrices belong to each of a finite number of states, grouping matrices based on a similarity metric. Since the dimensionality of an FC matrix with *N* nodes is *N*(*N* − 1)/2, and parcellations can have up to hundreds of nodes, clustering FC matrices into dFC states is a high-dimensional (unsupervised) learning task. Nevertheless, this step is usually performed using k-means clustering (Allen et al., 2014) (although other studies have used spectral clustering (Xu et al., 2018) or hierarchical methods (Ou et al., 2015; Yang et al., 2014)).

Several studies have evaluated the ability of SWC to detect dynamic changes in brain activity (Hindriks et al., 2016; Leonardi and Van De Ville, 2015; Lindquist et al., 2014; Thompson et al., 2018) and the effect of specific window parameters on its efficacy (Mokhtari et al., 2019; Savva et al., 2020; Shakil et al., 2016). However, fewer have quantitatively assessed the performance of the clustering step (Shakil et al., 2016). Some studies have provided validation of dFC clustering methods using synthetic (Allen et al., 2014; Erhardt et al., 2012; Lin et al., 2021; Mokhtari et al., 2019) or surrogate data (Shakil et al., 2016). These studies consistently demonstrate that k-means performance is highly dependent on sliding window shape and length (Lehmann et al., 2017; Mokhtari et al., 2019; Shakil et al., 2016) and on the signal-to-noise ratio (Lin et al., 2021; Shakil et al., 2016). Gonzalez-Castillo et al. (2015) used task-based fMRI to enforce switching between cognitive states, treating each task block (rest, memory, video, maths) as a different dFC state, therefore providing a “ground truth” in real-world data. They reported high accuracy when clustering at the subject level, and with a large number of regions in the parcellation (>100), but diminishing performance with smaller parcellations and shorter sliding window lengths.

K-means has been widely adopted for dFC analysis in cohort studies, to assess fractional occupancy and dwell time in a range of neurological disorders including Parkinson’s disease (Díez- Cirarda et al., 2018; Fiorenzato et al., 2019; Kim et al., 2017), schizophrenia (Bolton et al., 2020; Damaraju et al., 2014; Du et al., 2016, 2018; Fu et al., 2021; Rashid et al., 2014; Su et al., 2016), lewy body dementia (Schumacher et al., 2019) and autism (He et al., 2018a; Li et al., 2020; Rabany et al., 2019), in addition to healthy cognition (Hutchison and Morton, 2015) and sleep (Damaraju et al., 2020; Zhou et al., 2020). In applications such as these that compare the spatiotemporal dynamics of the brain across different groups, sources of heterogeneity between subjects, such as the shape of the hemodynamic response function (HRF) and levels of noise (Lehmann et al., 2017), may induce between-subject differences masking underlying dFC states. It is clear that inaccuracies in the clustering step would lead to inaccuracies in the measurement of properties such as fractional occupancy and dwell time, and therefore potentially spurious conclusions regarding the differences between groups.

It is therefore crucial to assess the ability of k-means to accurately quantify spatiotemporal dFC patterns and their transition statistics when applied to cohorts of multiple subjects. A particularly pertinent issue is that distance-based clustering methods like k-means do not perform well in high dimensional problems (Assent, 2012), and dFC analysis certainly fits into this category. In other applications of clustering to high-dimensional data, dimensionality reduction methods are often applied prior to clustering in order to map the data to a lower dimensional space. This provides salient features to the subsequent clustering step, reducing the effects of irrelevant or noisy features (Assent, 2012; Kriegel et al., 2009).

In this study, we evaluate the use of dimensionality reduction methods for feature selection prior to clustering dFC states (Fig. 1). We propose the use of deep autoencoders for feature selection prior to applying k-means clustering to the encoded data. We compare this deep clustering method to feature selection using principle component analysis (PCA), uniform manifold approximation and projection (UMAP), as well as applying k-means to the original feature space using either L1 or L2 distance. We provide extensive quantitative evaluation of clustering performance using synthetic datasets, representing data from multiple, heterogeneous artificial subjects (with subject-specific state time courses and noise parameters and variable intervals between state transitions). We measured performance in terms of clustering accuracy, similarity between the FC matrices of extracted states and those of true states, as well as error in measurements of fractional occupancy and dwell time. In synthetic data we find that deep clustering outperforms the other approaches. We then demonstrate the application of each method to real-world data from human subjects and show that the choice of feature selection method has a significant effect on group-level measurements of state temporal properties.

## 2. Methods

### 2.1. Data

We study both synthetic and real-world fMRI data. In general, data acquired for SWC analysis typically consist of several minutes of resting state fMRI data per subject. Nodes are defined either by a predetermined structural (Tzourio-Mazoyer et al., 2002) or functional (Craddock et al., 2012; Shen et al., 2013) parcellation scheme, or by generating a study-specific map by applying independent component analysis (ICA) at the group level (Kiviniemi et al., 2009). The timeseries of *T* time points by *N* nodes is then constructed by calculating the average BOLD signal within each region.

#### 2.1.1. Synthetic Data

We produced synthetic data using SimTB (Erhardt et al., 2012) (https://trendscenter.org/software/simtb/) to simulate BOLD activity in a set of *N* nodes under a model of spatiotemporal separability, using code modified from that originally used in Allen et al. (2014). In this model, timeseries data are constructed by linearly convolving a sequence of neural events with a HRF. We generated data governed by a time course of underlying dFC states, where the state occupied at a given time dictates the influence of each node’s activity on all other nodes, as follows. At each time point, a state-specific neural event occurs with some probability (set to the default value of 0.5). When a state-specific event occurs in a node, this has an additive or subtractive effect on the amplitude of events in all nodes which are functionally connected to this node, as defined by the dFC state occupied at that time step. In addition to these state-specific neural events, unique events occur randomly and are added to the time course for each node, representing spontaneous node-specific fluctuations in brain activity. For validation, the amplitude and probability of occurrence of these unique events was varied in order to create synthetic data with different noise levels.

We aimed to assess dFC clustering performance when applied to cohorts containing multiple subjects, so we created artificial subjects with simulated FC time courses. For each artificial subject, the underlying time course of dFC states was sampled from a hidden Markov model (HMM). Between-subject differences in dFC can be caused by individual differences in properties such as noise levels (i.e. neural noise and measurement noise) and HRF shape (Lehmann et al., 2017). Therefore, the probability and amplitude of unique neural events, the amplitude of the gaussian noise added to the BOLD signal, and the parameters of the HRF were varied across artificial subjects. The functionality to randomise HRF parameters is built into SimTB (Erhardt et al., 2012). The probability and amplitude of unique neural events and gaussian noise were sampled from normal distributions shown in Supplementary Table 1. We also aimed to ensure that clustering performance was independent of the state FC matrices and state transition matrix, so we grouped subjects into separate datasets. State FC matrices, and the transition matrix governing the HMM, were randomly generated in order to be unique to each dataset. Supplementary Fig. 1 summarises the structure of the synthetic datasets.

Each dataset was generated with a repetition time (i.e. the sampling rate), *T*_*R*_, of 2 s and an overall duration of 270 *T*_*R*_ (9 minutes). The method used to randomly generate the state FC matrices and transition matrix are described in the Supplementary Materials. Fig. 2a shows examples of randomly generated sets of dFC states. Fig. 2b shows examples of randomly generated transition matrices, with examples of corresponding state time courses shown in Fig. 2c. Note that we did not alter the functionality of the SimTB model, we simply automated the process of generating batches of synthetic data.

**Figure 2:**
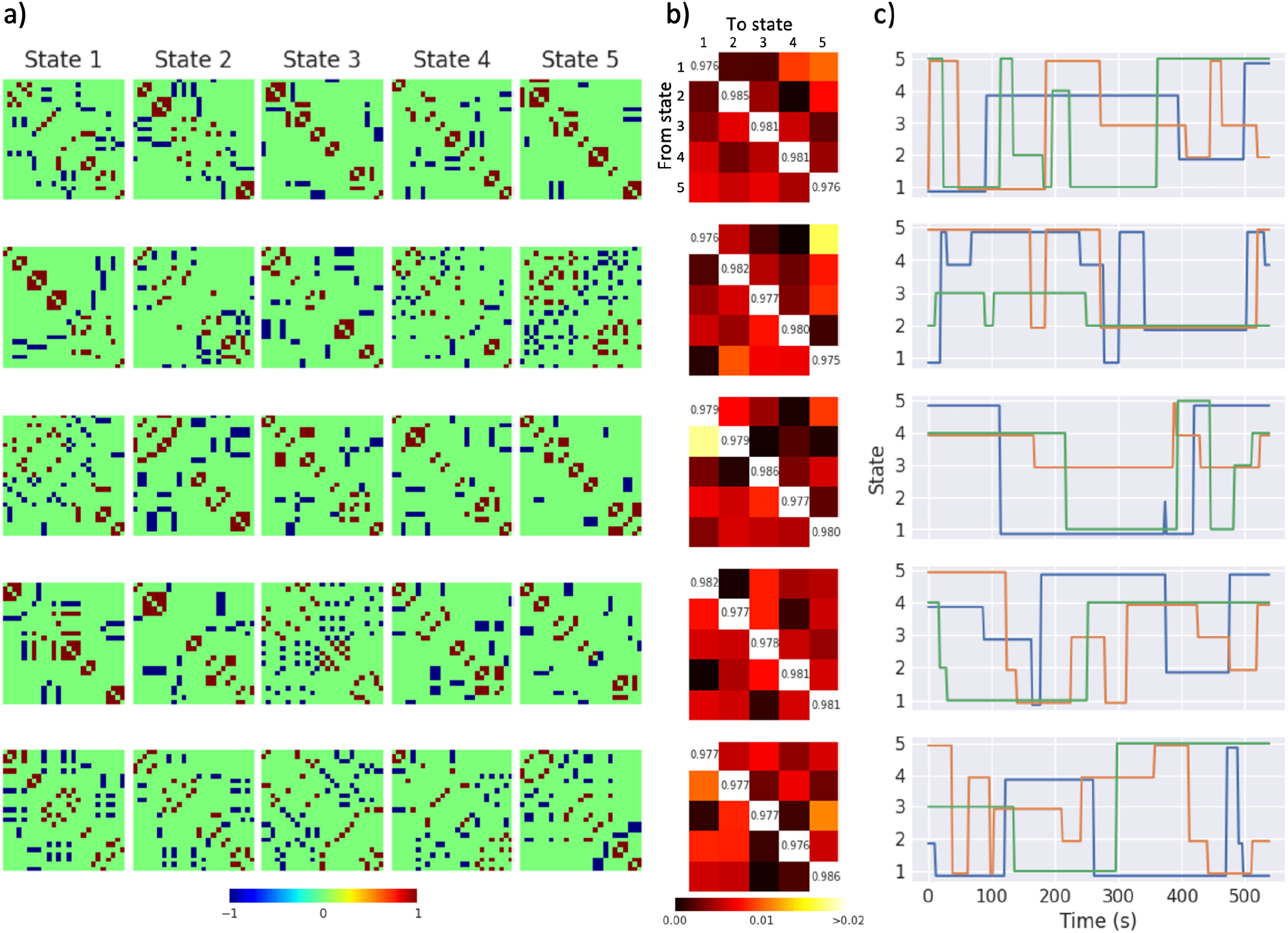
Examples of randomised states, transition matrices and time courses. a) Each row shows a set of five randomly generated dFC states, with functional connectivity indicated by the colour bar. b) Randomly generated transition matrices are shown, with the value in position (*i, j*) indicating the probability of switching from state *i* to state *j*, with probability indicated by the colour bar. For each transition matrix, the corresponding row in c) shows three examples of state time courses sampled from a HMM.

This model was used to generate training datasets to tune parameters of the dimensionality reduction methods (see Section 2.3), as well as validation datasets with different noise levels, number of nodes, number of states, number of subjects and HRF model in order to assess clustering performance in a range of experimental conditions (see Section 2.4.1).

#### 2.1.2. Human Data

For application to real-world data, we obtained resting-state fMRI data from the HCP1200 release from the Human Connectome Project (Van Essen et al., 2013) (https://www.humanconnectome.org). These data are provided as fully-processed subject-specific BOLD timeseries using a parcellation derived with spatial ICA. Briefly, processing steps which have already been applied to these data include spatial preprocessing according to Glasser et al. (2013) and temporal preprocessing according to Smith et al. (2013). Spatial preprocessing included correction for spatial distortions caused by gradient nonlinearity, correction for head motion, correction for *B*_0_ distortion global intensity normalisation, and 2 mm full-width at half maximum (FWHM) smoothing. Temporal preprocessed included high-pass temporal filtering (>2000s FWHM) and regression of artefact and motion-related time courses (Griffanti et al., 2014; Salimi-Khorshidi et al., 2014) followed by temporal demeaning and variance normalisation (Beckmann and Smith, 2004). Following preprocessing, spatial ICA was applied using MELODIC (Beckmann and Smith, 2004; Hyvarinen, 1999; Smith et al., 2014) from FSL (Smith et al., 2004), to obtain group-level parcellations. The set of ICA spatial maps was then mapped onto each subject’s BOLD timeseries to derive the node timeseries for each individual. We used the parcellation with *N* = 50 brain regions. We selected data which had no notable quality control issues recorded and used the first acquisition from each subject. These data had *T*_*R*_ = 720 ms and a duration of 1200 *T*_*R*_ (14 minutes 24 seconds).

### 2.2. Sliding-Window Correlations

The multivariate timeseries consisting of *T* time points and *N* brain regions were converted into a series of FC matrices using SWC, as follows. A window was used to select a short segment of the timeseries for all nodes. The window was then shifted in time by a given step size to extract overlapping segments, of the same length, for the whole timeseries of a given subject. In the synthetic data, we tested both rectangular and tapered (Hamming and Hanning) window shapes, and window lengths in the range 30–60 s (see Section 2.4.1) with a step size of 1 *T*_*R*_ (2 s). We measured FC in each window by estimating covariance from the precision matrix, regularised with the L1-norm (Allen et al., 2014; Smith et al., 2011; Varoquaux et al., 2010), where the regularisation parameter, λ_*L*1_, was estimated for each subject using cross-validation.

### 2.3. Clustering & Dimensionality Reduction

As the FC matrices are symmetric, the upper triangle was extracted and vectorised. Thus, after windowing and vectorisation, the BOLD data were transformed into an *X* by *Y* matrix, where *X* represents the number of subjects multiplied by the number of windows per subject, and *Y* represents the pair-wise correlations (equal to *N*(*N* − 1)/2). The clustering task that we focus on is then the assignment of each column of this data to a cluster.

We tested the performance of “raw” k-means against k-means applied after a dimensionality reduction step (Fig. 1). The dimensionality reduction procedures we used (described in detail below) are PCA, UMAP and deep clustering (autoencoder followed by k-means). The same k-means procedure was used in all methods. We compared our results to “chance” clustering by randomly assigning state labels to each window.

As the dimensionality reduction methods required parameter tuning, we generated training datasets with 50 subjects, 5 states, canonical HRF and medium noise, processed with a rectangular window of length 40 s. These data were used to tune the parameters of each method, using a grid search of parameter values to maximise clustering accuracy. A coarse grid of parameter values was used in order to prevent overfitting. Separate training datasets were generated with 15, 25 and 50 nodes, to re-tune clustering methods for the different input data dimensionality for these scenarios.

#### 2.3.1. k-means

We followed a k-means clustering methodology commonly used in dFC analysis (Allen et al., 2014). We selected exemplar FC windows at local maxima in variance and applied 128 repetitions of k-means (max 1000 iterations) to the FC matrices corresponding to these windows, each initialised with the k-means++ algorithm (Arthur and Vassilvitskii, 2006). The set of centroids which gave the lowest sum of squared error between each data point and its nearest centroid was then used to initialise k-means clustering (max 10000 iterations) for all windows. As well as the ‘default’ Euclidian (L2) distance metric, we also tested the Manhattan (L1) distance metric, as this is often used in dFC analysis due to high dimensionality (Aggarwal et al., 2001; Allen et al., 2014).

#### 2.3.2. Principle Component Analysis

PCA was applied to all FC matrices, then the first *p* principle components were used as features for k-means clustering, where *p* was chosen to maximise clustering accuracy using synthetic training data, as described above. The parameter values searched are shown in Supplementary Table 2. The resulting *p* for each parcellation is shown in Table 1.

**Table 1:** Parameters of each feature selection method for each size parcellation tested, as determined by a coarse grid search to maximise clustering accuracy using synthetic training data. The values searched are shown in Supplementary Tables 2–4. *p* is the number of principle components used for PCA. For UMAP, *m* is the number of neighbours used to determine the local connectivity of the high-dimensional graph before optimising the low-dimensional representation, *v* is the minimum permissible distance between points in the low-dimensional representation, and *u* is the number of dimensions. For deep clustering, *d*_1_, *d*_2_ and *d*_3_ are the number of units in the layers of the symmetric autoencoder (giving *d*_3_-dimensional encoded data).

| Number of nodes<br>N | Dimensionality<br>$\frac{N(N-1)}{2}$ | PCA<br>$p$ | UMAP<br>$m$ $v$ $u$ | | | Deep Clustering<br>$d_1$ $d_2$ $d_3$ | | |
| --- | --- | --- | --- | --- | --- | --- | --- | --- |
| 15 | 105 | 16 | 30 | 1.0 | 32 | 512 | 256 | 16 |
| 25 | 300 | 32 | 30 | 1.0 | 64 | 512 | 256 | 32 |
| 50 | 1225 | 256 | 40 | 1.0 | 32 | 1024 | 256 | 64 |

#### 2.3.3. Uniform Manifold Approximation and Projection

UMAP is a nonlinear dimension reduction technique which projects data onto a low-dimensional manifold by constructing a high-dimensional graph representation, then optimising a low-dimensional graph to have a structure as similar as possible to the high-dimensional graph (McInnes et al., 2018). The structure of the high-dimensional graph is determined locally based on distances to the nearest *m* neighbours. Higher *m* results in a low-dimensional projection which more accurately captures the global structure of the data rather than preserving local distances to neighbours. Additional parameters which must be chosen are the number of dimensions, *u*, in the low-dimensional subspace, and the minimum permissible distance, *v*, between points in the low-dimensional representation. We used UMAP to embed all FC matrices into a low-dimensional subspace, then applied k-means clustering to the embedded data. The values of *u, v* and *m* were chosen to maximise clustering accuracy using synthetic training data, as described above. The parameter values searched are shown in Supplementary Table 3. The tuned parameters for each size parcellation are shown in Table 1.

#### 2.3.4. Deep Clustering

Deep learning has provided powerful tools for neuroimaging analysis, including segmentation of anatomical structures or lesions in structural MRI (Akkus et al., 2017), annotation of cognitive states in task-based fMRI data (Zhang et al., 2021), or clinical diagnosis from functional connectivity networks (He et al., 2018b; Kam et al., 2019; Vieira et al., 2017; Wang et al., 2020) (for an overview of deep learning concepts and methodology, and a review of applications to studies of neurological disorders, see Vieira et al. (2017)). Whereas these supervised applications of deep learning require a large amount of ground truth data for training (Quaak et al., 2021), autoencoders can be used as a feature selection method for unsupervised applications.

Autoencoders are a type of artificial neural network which copy the input data to the output, via a low-dimensional encoding layer (Goodfellow et al., 2016; Vincent et al., 2008). The bottleneck formed by this low-dimensional encoding layer forces the network to extract features from which the original data can be reproduced via the decoding layers. In this case, the input data is used as the training target, thus autoencoders can be used for feature selection or dimensionality reduction in unsupervised clustering applications with no ground truth (Guo et al., 2017; Xie et al., 2016). Here, we use autoencoders as a data-driven approach to determining feature space at the group level, allowing clustering to be applied to the salient features provided by the low-dimensional encoding layer. This framework is known as deep clustering (Caron et al., 2018; Guo et al., 2017). The proposed deep clustering framework consists of training an autoencoder on all FC windows before applying k-means clustering to the encoded data (Fig. 3). We used a fully-connected autoencoder with three encoding layers and a symmetric decoder. Weights were trained using the Adam optimiser (Kingma and Ba, 2014) to minimise the mean-squared error (MSE) between the input and output, trained for 100 epochs with a batch size of 50. Rectified linear unit (ReLU) activation functions were used for hidden layers and linear activation functions were used for the low-dimensional layer and output layer. The number of units in each layer were chosen to maximise clustering accuracy using synthetic training data, as described above. The parameter values searched are shown in Supplementary Table 4. The number of units in each layer for each parcellation is shown in Table 1.

**Figure 3:**
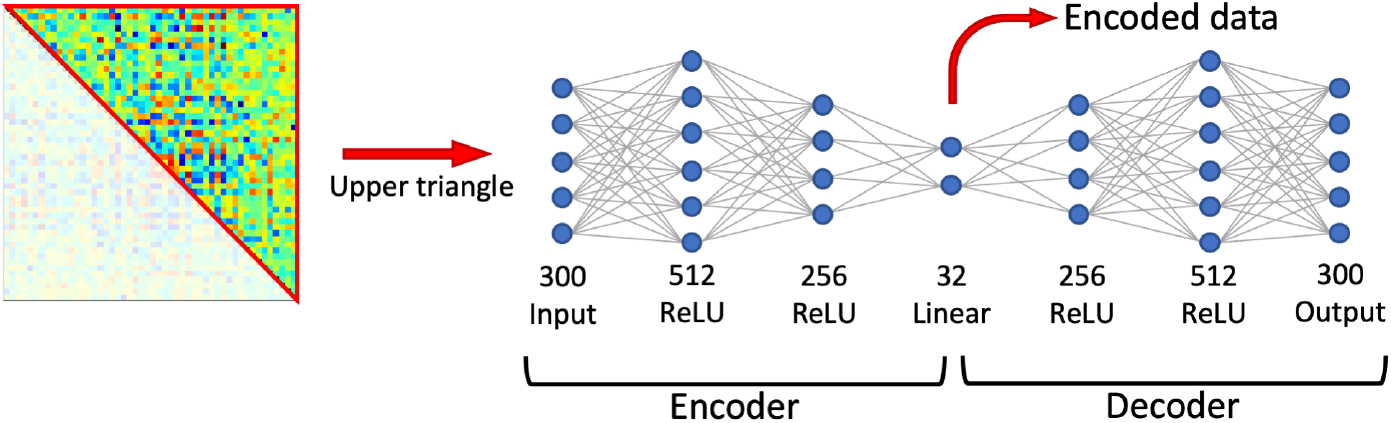
Autoencoder architecture. The architecture, shown here for a parcellation with *N* = 25 nodes, was determined by a coarse grid search to maximise clustering accuracy using a synthetic training dataset. The upper triangle of each FC matrix is taken as an input. Each layer shows the number of units and the activation function. The encoded data is used for clustering. ReLU = rectified linear unit.

### 2.4. Experiments

#### 2.4.1. Clustering Synthetic Data

We varied the parameters of the model and preprocessing steps to assess clustering performance in each of the following cases:

a. Number of “subjects”: 10, 50 and 100.
b. Number of regions in the parcellation: 15, 25 and 50.
c. Number of states: 3, 5 and 7.
d. Hemodynamic response function (HRF): Both HRF models provided in SimTB were tested; the canonical HRF, and the Windkessel-Balloon model.
e. Noise: Low, medium and high noise datasets were generated by varying the probability and amplitude of unique events in the underlying neural time course, and the amplitude of Gaussian noise added to the BOLD signal. The distributions of these parameters are shown in Supplementary Table 1.
f. Sliding window shape: Rectangle, Hamming and Hanning.
g. Sliding window length: 30, 40 and 60 s.

In each dataset, parameters that were not varied were set to the following default values: 50 subjects; 25 nodes; canonical HRF; high noise; rectangle sliding window length of 40 s. For validation, clustered states were matched to true states by pairing those with maximum cosine similarity between corresponding FC matrices. To allow comparison of dFC state centroids between clustering methods, the representative FC matrix for a given state was constructed by averaging all FC windows belonging to that state (rather than using the low-dimensional centroid derived by the k-means step). Clustering performance was then measured by accuracy (fraction of correctly labelled windows), dwell time MSE, fractional occupancy MSE, and the mean cosine similarity between cluster centroids and true states. To calculate dwell time MSE and fractional occupancy MSE, these properties were calculated for each state in every subject, then the squared error between these measurements and the true values were calculated and averaged across subjects and states. For each parameter set, we constructed five datasets (each with a unique set of state FC matrices and a unique transition matrix) and performed 10 runs of each method on each dataset. We then averaged performance metrics over all 50 runs of each method on the given parameter set.

To demonstrate the differences between measurements of temporal properties across subjects in each dataset, we took results from one run of each method and performed an unpaired two-tailed t-test to test for significant differences from the true distribution of fractional occupancy and dwell time measurements for each state.

#### 2.4.2. Clustering Real-World Data

We selected five non-overlapping groups of 100 subjects from the Human Connectome Project dataset (see Section 2.1.2). In each group, we applied SWC with a rectangular window of length 55 *T*_*R*_ (39.6 s) with a step size of 2 *T*_*R*_ (1.44 s), giving 573 windows per subject. This was based on previous work suggesting that, where motion noise is not excessive, a rectangular window is suitable for detecting dFC (Savva et al., 2020). The number of clusters, *k*, was selected for each dataset using the elbow criterion of the within-cluster to between-cluster distance when clustering exemplar FC windows (Allen et al., 2014). In all groups of 100 subjects, we identified the optimal number of clusters to be *k* = 4. We then applied clustering with each method to each group, using the parameters shown in Table 1 for 50 nodes. For subsequent comparison, states were matched between methods by pairing those with maximum cosine similarity between the corresponding FC matrices.

To assess the measurements of state temporal properties provided by each method, we calculated the fractional occupancy and dwell time of each state in each subject. To determine whether the stochasticity introduced by training the autoencoder affected measurements of state temporal properties across repeated runs of deep clustering, we performed 10 runs of deep clustering on each dataset and used a one-way analysis of variance (ANOVA) to test for differences between runs in the fractional occupancy and dwell time of each state.

To determine whether the choice of feature selection method affected the measurement of state temporal properties, we performed one run of each method and used a one-way ANOVA to test for differences between methods in fractional occupancy and dwell time. False discovery rate (FDR) correction was applied using the Benjamini-Hochberg method (Benjamini and Hochberg, 1995). If this yielded significant results (FDR-corrected *p* < 0.05) then post hoc pairwise comparison between methods was performed using unpaired two-tailed t-tests with FDR correction applied.

### 2.5. Data Availability Statement

Real-world fMRI data were obtained from the Human Connectome Project (Van Essen et al., 2013) (https://www.humanconnectome.org). The SimTB model (Erhardt et al., 2012) used to generate synthetic data was obtained from https://trendscenter.org/software/simtb/. The code used in this study is publicly available at https://github.com/apcspencer/dFC_DimReduction.

## 3. Results

### 3.1. Validation

To examine the efficacy of each dimension reduction method, we quantified clustering performance using synthetic data. Of the methods tested, deep clustering gave the highest accuracy in all synthetic datasets overall (Fig. 4). Deep clustering provided the highest mean cosine similarity between extracted states and true states, the lowest MSE in fractional occupancy, and the lowest MSE in dwell time in the majority of datasets (Fig. 4). The only exceptions are those with 10 subjects, 15 nodes and 30s windows, where PCA and UMAP give marginally better results for mean cosine similarity, and lower MSE in dwell time, respectively. Additionally, deep clustering was the only method to give measurements of fractional occupancy and dwell time better than chance in all datasets.

**Figure 4:**
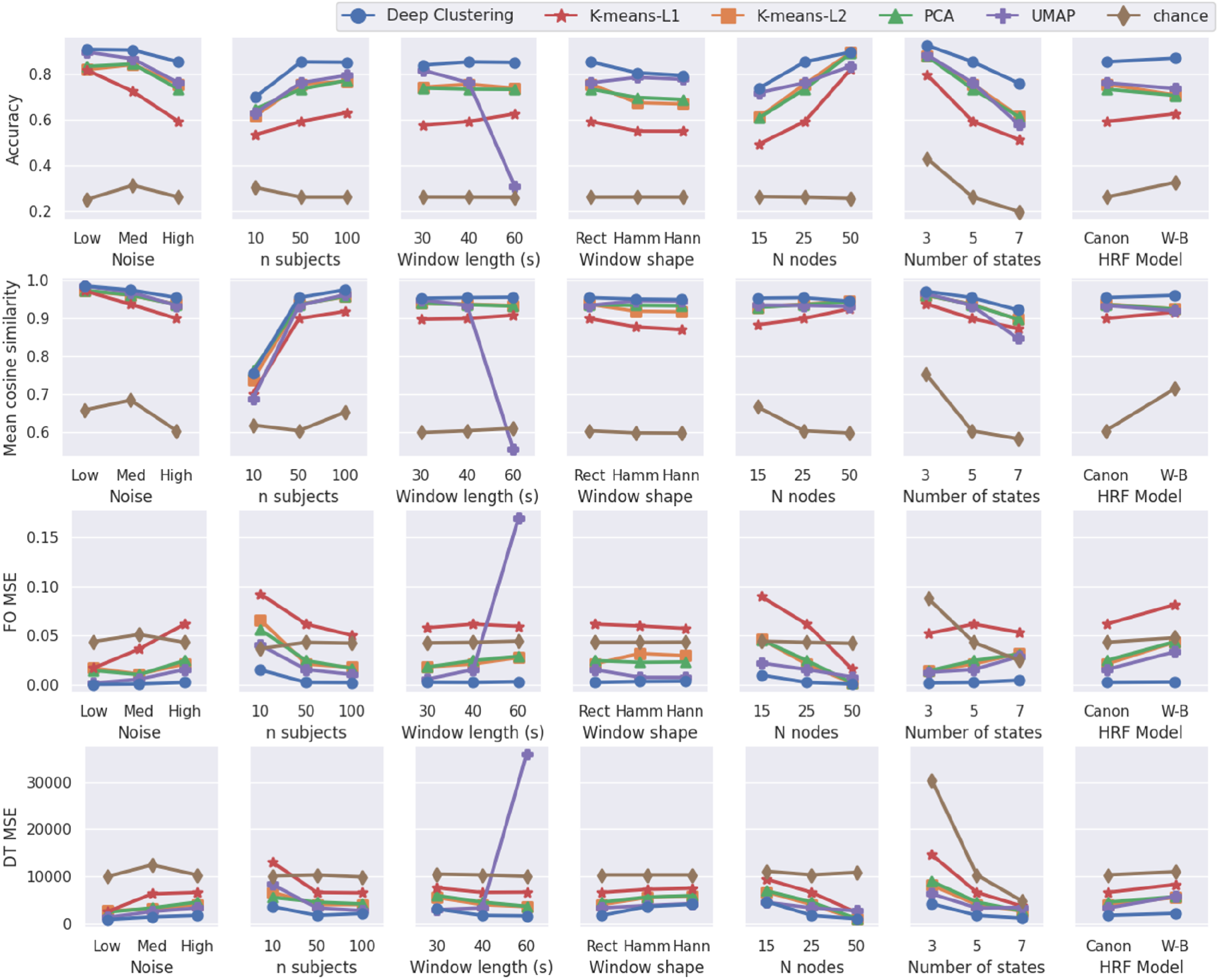
Results of validation with synthetic data. Performance metrics from each feature selection method plotted for synthetic validation datasets with varying model parameters. For each set of model parameters, 5 datasets were generated with the given parameters, each with a unique transition matrix and set of state FC matrices, then 10 runs of each method were applied to each of these datasets. The performance averaged over these 50 runs is plotted for each parameters set. The accuracy of each individual run is shown in Supplementary Fig. 2. FO = fractional occupancy; DT = dwell time; MSE = mean squared error; PCA = principle component analysis; UMAP = uniform manifold approximation and projection; HRF = hemodynamic response function; Rect = rectangular; Hamm = Hamming; Hann = Hanning; Canon = canonical; W-B = Windkessel-Balloon.

Applying k-means to the original feature space, using L1 distance almost always gave the worst performance. In most datasets, using PCA for feature selection prior to k-means did not offer any performance improvements over applying k-means with L2 distance to the original feature space. UMAP gave variable accuracy, with good performance in data with low noise, 30 s window, Hamming window, or 15 nodes, but performed poorly in the 60 s window and 7 state datasets, in which fractional occupancy measurements were worse than chance.

The accuracy of each individual run of each method, grouped by dataset, is shown in Supplementary Fig. 2. Repeated runs of each method on a given dataset had tightly grouped accuracy scores, with the exception of datasets in the 10 subject parameter set, which was highly variable. In terms of accuracy, the ranking of the methods was similar all five datasets for each set of parameter values, with deep clustering almost always performing best, and k-means using L1 distance almost always performing worst.

Fig. 5 shows exemplar data from the results of each clustering algorithm applied to a high noise dataset. It can be seen that the distribution of dwell time measurements for deep clustering were not statistically distinguishable from the ground truth. On the other hand, these were significantly different from the ground truth for states 1, 2 and 5 with PCA or k-means using L1 distance, and for state 1 with k-means using L2 distance (Fig. 5c). Additionally, fractional occupancy measurements from k-means using either L1 or L2 distance, PCA and UMAP were significantly different from the truth for state 3. The high fractional occupancy measurements for state 3 show that a large number of FC windows were incorrectly assigned to this cluster. This has a visible effect on the state FC matrices for state 3 for these methods (Fig. 5a), in which most elements are close to zero due to being averaged over a large number of FC windows. Conversely, all five state FC matrices derived using deep clustering have visually similar structure to the ground truth.

**Figure 5:**
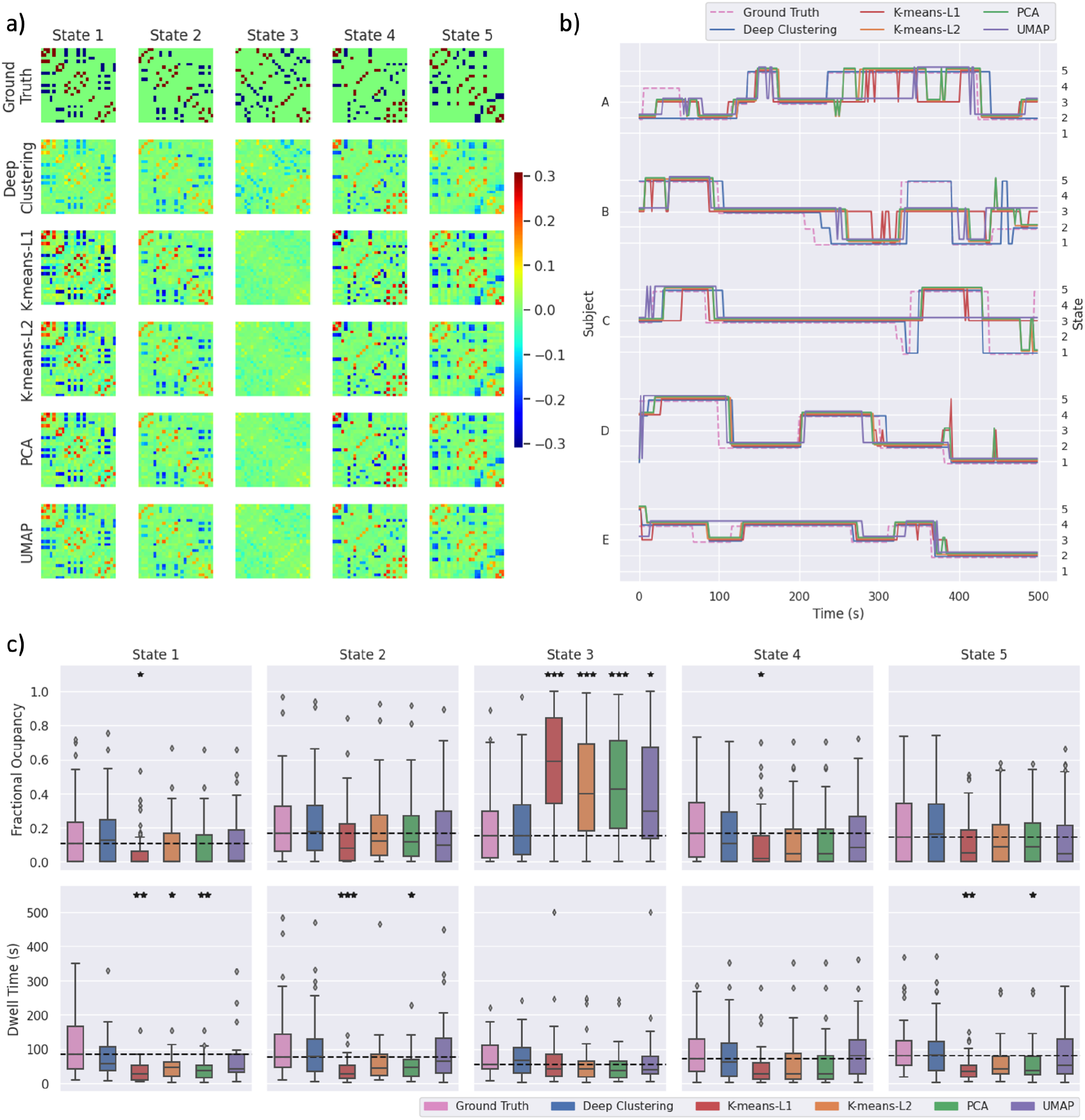
Clustering results from one run of each feature selection method applied to synthetic data with high noise. a) State FC matrices are plotted with connectivity indicated by the colour bar. b) State time courses are shown for five subjects. c) The distribution across subjects of fractional occupancy and dwell time measurements are plotted for each state. Boxes show the interquartile range, with a line for the median. The median of the ground truth is shown as a dashed line across each plot for comparison with other methods. Whiskers extend to the range of the data, not including outliers which are shown as diamonds. Significant differences from the true distributions, measured by unpaired two-tailed t-tests, are indicated as follows: \**p* < 0.01, \*\**p* < 0.001, \*\*\**p* < 0.0001.

A similar effect is observed with most parameter sets (see Supplementary Fig. 3–15), which clearly demonstrates the difference in performance between deep clustering and the other methods. K-means using either L1 or L2 distance, PCA and UMAP derive state FC matrices that appear to be very similar to the ground truth for most states (see e.g. states 1, 2, 4 and 5 in Fig. 5a). It appears this is because these methods only identify the FC windows that very strongly express the state’s pattern of connectivity. Spurious correlations in many other FC windows cause those windows to be incorrectly assigned to a single state by these methods (state 3 in Fig. 5a). We postulate that for deep clustering, the autoencoder projects the data to a low-dimensional feature space in which the salient features allow the FC windows to be assigned to the correct state more accurately, reducing the effect of the spurious correlations.

Notably, PCA as well as k-means using either L1 or L2 distance each gave lower measurements of median dwell time than the ground truth for all states. This was the case for all parameter sets (see Supplementary Fig. 3–15). This is likely due to occasional spurious switching between states, as seen in the state time courses of individual subjects plotted in Fig. 5b.

### 3.2. Application to Real-World Data

We applied each method to real-world data from five groups of 100 subjects from the Human Connectome Project. Clustering results for the first group are shown in Fig. 6. The estimated state FC matrices were largely comparable for all methods except for UMAP, though state 4 also differed for deep clustering compared to the other methods (Fig. 6a). There were significant differences in the measurements of fractional occupancy for states 1 and 3, and dwell time in states 2–4 (Fig. 6b; one-way ANOVA, FDR-corrected *p* < 0.05).

**Figure 6:**
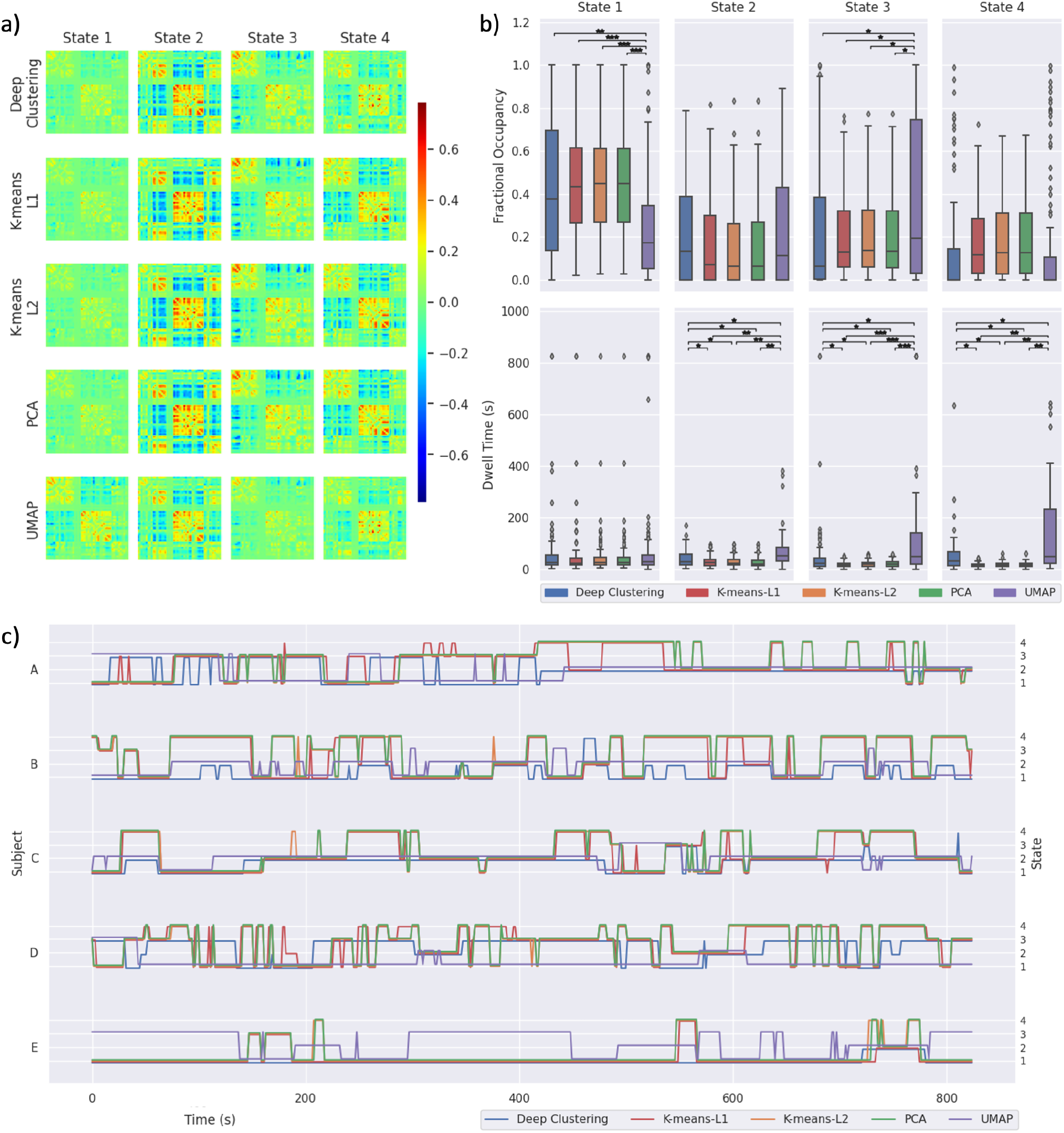
Human Connectome Project clustering results from each feature selection method. a) State FC matrices resulting from each method, with connectivity indicated by the colour bar. b) Fractional occupancy and dwell time measurements across subjects are shown for each state. Boxes show the interquartile range with a line for the median. Whiskers extend to the range of the data, not including outliers which are shown as diamonds. Measurements were compared using a one-way ANOVA, with post hoc pairwise comparison using two-tailed t-tests. c) State time courses are shown for five subjects. FDR-corrected: \**p* < 0.05, \*\**p* < 0.001, \*\*\**p* < 0.0001.

Results for the four additional groups of 100 subjects are shown in Supplementary Fig. 16–19. Notably, PCA and k-means using either L1 or L2 distance resulted in lower dwell time measurements than deep clustering and UMAP for most states in all groups, which is similar to our finding when analysing synthetic data (see Fig. 5 and Supplementary Fig. 3–15). State time courses are plotted for five subjects, demonstrating qualitative differences in the switching characteristics of each method (Fig. 6c).

To assess whether the stochasticity introduced when training the autoencoder affected measurements of state temporal properties across repeated runs of deep clustering, we performed a one-way ANOVA to test for differences in the fractional occupancy and dwell time of each state for 10 repetitions applied to each group of 100 participants. Measurements of both fractional occupancy and dwell time were robust across repeated runs for all states in all groups of participants (lowest uncorrected *p*-value = 0.1191). These results are shown in full in Supplementary Fig. 20.

## 4. Discussion

In this study, we evaluated the use of dimensionality reduction methods prior to clustering dFC data from SWC. We proposed a deep clustering framework consisting of training a fully-connected autoencoder on all FC windows, followed by applying k-means clustering to the encoded data. We quantitatively assessed clustering performance using multiple synthetic datasets, each with unique sets of state FC matrices and transition probabilities, and each containing multiple timeseries representing different subjects with randomised model parameters. We demonstrated that deep clustering gives the highest accuracy across a range of model parameters and preprocessing conditions, including varying HRF parameters, noise levels (both additive Gaussian noise and spurious neural events), number of subjects, number of states, number of nodes, and sliding window shape and length. Further, when measuring the error in estimates of fractional occupancy and dwell time, deep clustering was the only method to perform better than chance across all experimental conditions.

PCA and both k-means approaches consistently performed worse than deep clustering, with k-means using L1 distance usually performing worst, and PCA or k-means using L2 distance giving roughly similar performance. UMAP gave good performance in some tests, with similar accuracy to deep clustering in the low noise datasets, 30 s window datasets, and 15 node datasets. However, the performance of UMAP varied widely, giving comparatively lower accuracy in the 7 state datasets or the datasets with the Windkessel-Balloon HRF, and gave very poor performance in the 60 s window datasets. This is likely due to the fact that UMAP relies on distances between data points (windowed FC matrices in this case) to construct the low-dimensional feature space. As such, UMAP parameters are unlikely to generalise to data in which distances between data points, and the number of data points, are drastically different from the tuning data, which may be the case for a different window size. As we can’t tune the feature selection methods on real-world data, these different synthetic validation datasets serve as an indication of how well the performance of a given method generalises, and therefore its reliability when applied to real-world data which may be much further from the tuning dataset. As UMAP gives variable performance, declining dramatically in some cases, it is clearly not robust across parameters and is therefore not the most reliable for application to real-world data.

When applied to real-world data, measurements of state temporal properties were dependent on the choice of feature selection method. For many states, PCA and both k-means approaches (using either L1 or L2 distance) gave estimates of dwell time that were significantly lower than those given by deep clustering. This echos the results from synthetic data, in which these methods consistently underestimated dwell time. UMAP gave longer estimates of dwell time than other methods. This is likely due to the neighbour embedding algorithm placing consecutive windows, with high similarity, close to each other in the embedded space. This may not have been an issue in the simulated data which has sharper transitions between states (Shakil et al., 2016).

Measurements of state temporal properties in real-world data using deep clustering were consistent across repeated evaluations of the algorithm. While this reproducibility does not necessarily indicate good or meaningful clustering, it does show that deep clustering gives robust results despite the stochasticity introduced by the deep learning approach.

There are an increasing number of available approaches for dFC analysis of resting state fMRI data, including methods of extracting brain states directly from voxelwise BOLD data (Lin et al., 2021), or methods to determine spatially and temporally overlapping states (Karahanoğlu and Van De Ville, 2015) (for a review, see Preti et al. (2017)). However, applying k-means to SWC data is the most common approach in studies of dFC in neurological disorders, despite evidence showing that this method gives poor characterisation of state transitions (Shakil et al., 2016). Our results show that applying k-means to the original feature space, without the use of feature selection, may be insufficient for measuring state temporal properties, despite a high similarity between the FC matrices representing extracted states and true states. Our proposed deep clustering approach provides accurate measurements of temporal properties in synthetic data, robust to variations in model parameters and experimental conditions.

In neuroimaging studies, autoencoders have been used for feature selection prior to modelling (Suk et al., 2016), and for pre-training layers of a classifier (Heinsfeld et al., 2018). A recent study has applied autoencoders directly to fMRI BOLD data in order to improve individual identifiability of functional connectomes (Cai et al., 2021). The goals of Cai et al. (2021) were to residualise the BOLD data, by applying autoencoders directly to the BOLD timeseries to remove common neural activities, in order to enhance inter-subject variability for functional connectome ‘fingerprinting’. Conversely, we used autoencoders as a data-driven approach to determine feature space at the group level in order to improve clustering of FC windows, thus improving measurements of dFC state temporal properties. Our deep clustering approach is applied to the FC matrices, such that it fits into the existing, well-established framework of SWC analysis.

The use of deep learning often raises concerns of overfitting, as the number of parameters (weights in the neural network) exceeds the number of data points (FC windows) resulting in poor generalisation to data points outside the training set (Vieira et al., 2017). In the deep clustering approach used here, the autoencoder is always trained on the data being clustered, therefore the weights do not have to generalise to unseen data. The only parameters which are not trained on each data point encountered are hyperparameters such as the number of units in each layer, the activation function, the batch size and the number of epochs. By demonstrating that deep clustering performance is robust to model parameters and preprocessing parameters, we have shown that the autoencoder architecture and hyperparameters, which were tuned with a coarse grid search to maximise clustering accuracy using a synthetic training dataset, offer good generalisation.

The use of tapered sliding windows when computing SWC has been proposed in order to diminish the effect of spurious fluctuations causing large discontinuities when entering and leaving the window (Allen et al., 2014; Mokhtari et al., 2019; Shakil et al., 2016). Our results in synthetic data showed slightly worse clustering performance with all methods when using tapered windows, in comparison with a rectangular window. However, as previous suggested (Shakil et al., 2016), this is likely due to the sharp discontinuities at state transitions in synthetic data, which may not reflect the characteristics of state changes in real-world data. Additionally, we did not match the different window types (e.g. by varying window length to match the cuttoff frequency) (Mokhtari et al., 2019). The purpose of assessing different window shapes in this study was to evaluate clustering performance across a range of experimental conditions; a comprehensive comparison of sliding window parameters and preprocessing options is beyond the scope of this work, but can be found in previous studies (Hindriks et al., 2016; Leonardi and Van De Ville, 2015; Mokhtari et al., 2019; Shakil et al., 2016).

There is still some debate over whether dFC analysis truly captures underlying neural activity (Gonzalez-Castillo et al., 2015; Handwerker et al., 2012; Matsui et al., 2019) or simply artefacts due to head motion (Laumann et al., 2017). The improved performance offered by deep clustering may allow more refined assessment of the ability of SWC to capture neural activity.

## 5. Conclusion

We have demonstrated that a deep clustering framework, comprising autoencoders for feature selection prior to k-means clustering, offers improved dFC clustering performance on synthetic data. When applied to real-world data, this performance increase resulted in significant differences in the measurement of temporal characteristics of brain states compared to the standard approach. These differences qualitatively reflected the differences observed in synthetic clustering results.

## Acknowledgements

Data were provided in part by the Human Connectome Project, WU-Minn Consortium (Principal Investigators: David Van Essen and Kamil Ugurbil; 1U54MH091657) funded by the 16 NIH Institutes and Centers that support the NIH Blueprint for Neuroscience Research; and by the McDonnell Center for Systems Neuroscience at Washington University.

## Funding

AS is supported by the Wellcome Trust (WT220070/Z/20/Z).

## Declarations of interest

none.

## Notes

### Competing Interest Statement

The authors have declared no competing interest.

