## Supplementary Materials for "Using Deep Clustering to Improve fMRI Dynamic Functional Connectivity Analysis"

### Generating State FC Matrices and Transition Matrices

To generate each FC matrix, we chose the number of modules it should comprise a priori. We set the maximum module number to 5, 7 and 9 when generating data with 15, 25 and 50 nodes respectively (increased from 3 modules used when generating data with 10 nodes in the original code (Allen et al., 2014)). Then, each node was randomly assigned a sign (+/-) and a module number, with uniform probability. The FC matrix was then constructed by assigning a positive connection where nodes belonged to the same module and had the same sign, and a negative connection between nodes which belonged to the same module and had different sign. We then re-ordered the nodes such that the matrix had optimal community structure, using the *modularity_und* function from the Brain Connectivity Toolbox (`https://sites.google.com/site/bctnet/`) (Rubinov and Sporns, 2010), in order to produce dFC states with a visible modular structure to allow qualitative comparison of state FC matrices. This process was repeated for each state.

To generate the transition matrix governing the HMM, the probability of remaining in the same state $s$ was initially set to a value $p_s$ (i.e. values on the leading diagonal of the transition matrix). The remaining $1 - p_s$ was then randomly split between all states (including state $s$ in order to induce different dwell times in different states). The value of $p_s$ was determined by experimenting with values close to 1, in order to give a median switching timescale of around 60 s, and was varied for datasets with different number of states. We set $p_s = 1 - 0.005 \times n$, where $n$ is the number of states.


| Noise level | Unique event probability | Unique event amplitude | Gaussian noise amplitude |
|---|---|---|---|
| Low | $p_u \sim \mathcal{N}(0.3, 0.2^2)$ | $a_u \sim \mathcal{N}(0.3, 0.2^2)$ | $a_g \sim \mathcal{N}(0.3, 0.2^2)$ |
| Medium | $p_u \sim \mathcal{N}(0.5, 0.3^2)$ | $a_u \sim \mathcal{N}(0.5, 0.3^2)$ | $a_g \sim \mathcal{N}(0.4, 0.2^2)$ |
| High | $p_u \sim \mathcal{N}(0.6, 0.2^2)$ | $a_u \sim \mathcal{N}(0.7, 0.2^2)$ | $a_g \sim \mathcal{N}(0.6, 0.2^2)$ |

Supplementary Table 1: Parameters used to generate synthetic data with low, medium and high noise levels. In order to create heterogeneous subjects, each parameter was sampled from a normal distribution for each subject. The distributions used are shown for the unique event probability, $p_u$, unique event amplitude, $a_u$, and Gaussian noise amplitude, $a_g$, each displayed in the form $\mathcal{N}(\mu, \sigma^2)$ with a mean $\mu$ and standard deviation $\sigma$. See the main text for explanation of these parameters in the SimTB model.

| Nodes | Dimensionality | $p$ |
|---|---|---|
| 15 | 105 | [**16**, 32, 64] |
| 25 | 300 | [16, **32**, 64, 128] |
| 50 | 1225 | [16, 32, 64, 128, **256**, 512] |

Supplementary Table 2: Parameter values searched when tuning the number of principle components, $p$, to be used for clustering following dimensionality reduction with PCA. For each number of nodes in the parcellation, the dimensionality of the dFC matrices is shown, as well as the range of parameters searched during tuning. The values which gave maximal clustering accuracy in synthetic training data are shown in bold for each number of nodes.

| Nodes | Dimensionality | $m$ | $v$ | $u$ |
|---|---|---|---|---|
| 15 | 105 | [**30**, 40, 50, 60] | [0.1, 0.2, 0.5, **1.0**] | [**32**, 64] |
| 25 | 300 | [**30**, 40, 50, 60] | [0.1, 0.2, 0.5, **1.0**] | [32, **64**] |
| 50 | 1225 | [30, **40**, 50, 60] | [0.1, 0.2, 0.5, **1.0**] | [**32**, 64, 128] |

Supplementary Table 3: Parameter values searched when tuning the UMAP algorithm for dimensionality reduction. A grid search was performed over all combinations of values for these three parameters. $m$ is the number of neighbours used to determine the local connectivity of the high-dimensional graph before optimising the low-dimensional representation, $v$ is the minimum permissible distance between points in the low-dimensional representation, and $u$ is the number of dimensions. See the main text for explanation of these parameters. For each number of nodes in the parcellation, the dimensionality of the dFC matrices is shown, as well as the range of parameters searched during tuning. The values which gave maximal clustering accuracy in synthetic training data are shown in bold for each number of nodes.

| Nodes | Dimensionality | $d_1$ | $d_2$ | $d_3$ |
|---|---|---|---|---|
| 15 | 105 | [128, 256, **512**] | [64, 128, **256**] | [**16**, 32, 64] |
| 25 | 300 | [128, 256, **512**] | [64, 128, **256**] | [16, **32**, 64] |
| 50 | 1225 | [256, 512, **1024**] | [128, **256**, 512] | [16, 32, **64**] |

Supplementary Table 4: Parameter values searched when tuning the autoencoder architecture for deep clustering. A grid search was performed over all combinations of values for these three parameters. $d_1$, $d_2$ and $d_3$ are the number of units in the layers of the symmetric autoencoder (giving $d_3$-dimensional encoded data). For each number of nodes in the parcellation, the dimensionality of the dFC matrices is shown, as well as the range of parameters searched during tuning. The values which gave maximal clustering accuracy in synthetic training data are shown in bold for each number of nodes.

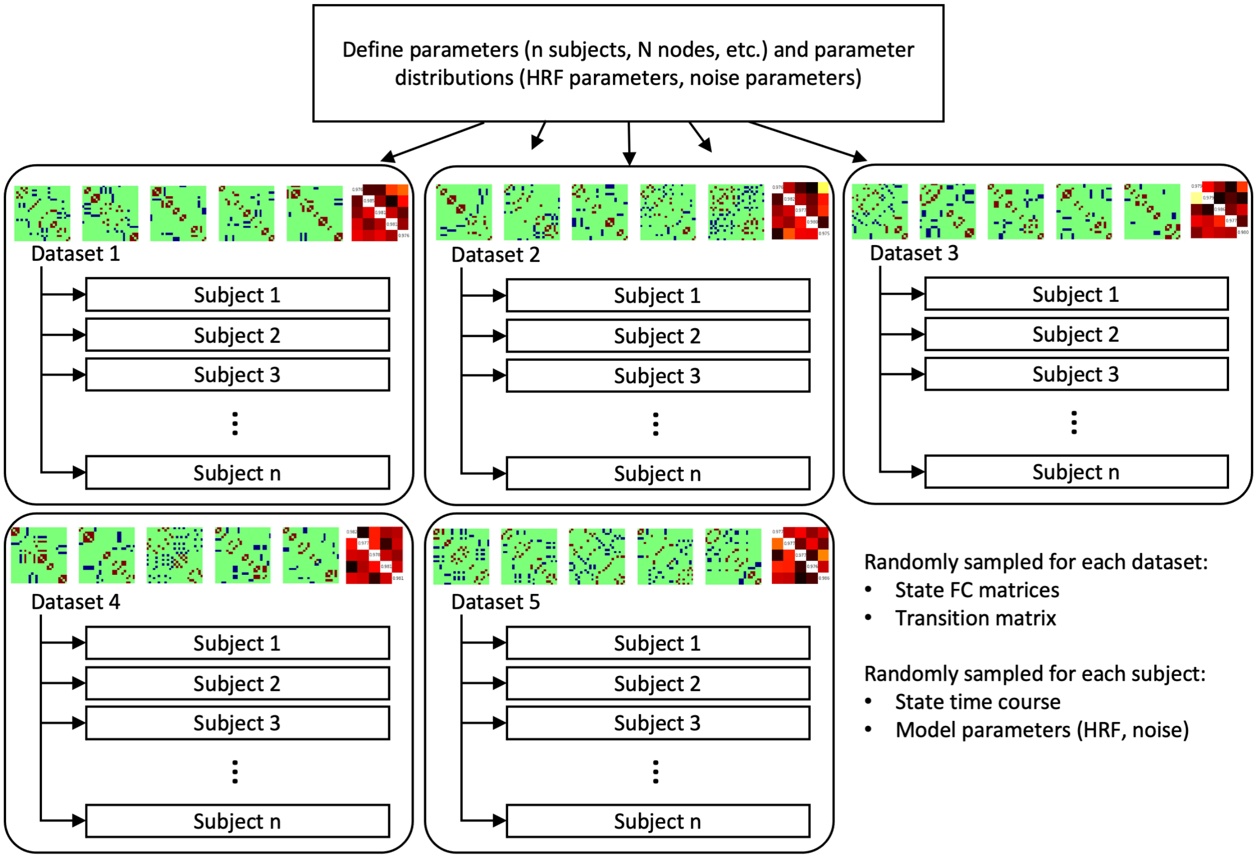


Supplementary Figure 1: Model hierarchy for generating synthetic datasets. First the parameter set is defined (including the number of subjects, number of nodes and distributions for noise and HRF parameters), then datasets can be generated, each with a unique set of state FC matrices and a unique transition matrix. Each dataset contains multiple subjects, which each have a randomised time course of underlying states, sampled from a HMM governed by the transition matrix, and HRF and noise parameters sampled from the predefined normal distributions. HMM = hidden Markov Model; HRF = hemodynamic response function.

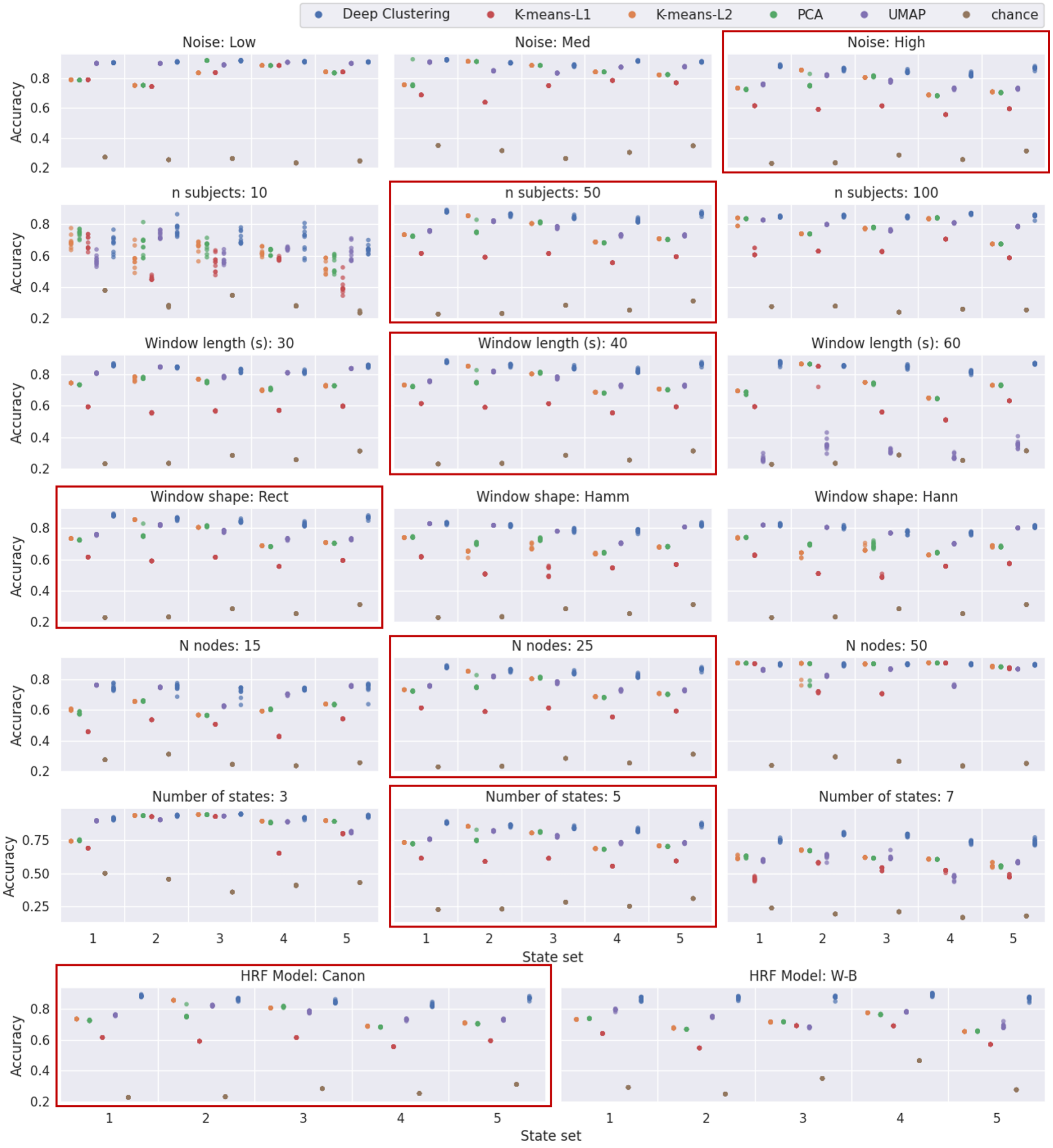


Supplementary Figure 2: Clustering accuracy in synthetic data. For each parameter set, five unique sets of states were randomly generated, each with a corresponding randomly generated transition matrix. Each method was applied to each dataset 10 times. The accuracy is plotted for each run of each method on each set of states with each parameter set. The red boxes indicate the default parameter sets, thus each of these plots show the same data, but are replicated to allow comparison with adjacent plots in which parameters are varied. Rect = rectangular; Hamm = Hamming; Hann = Hanning; HRF = haemodynamic response function; W-B = Windkessel-Balloon.

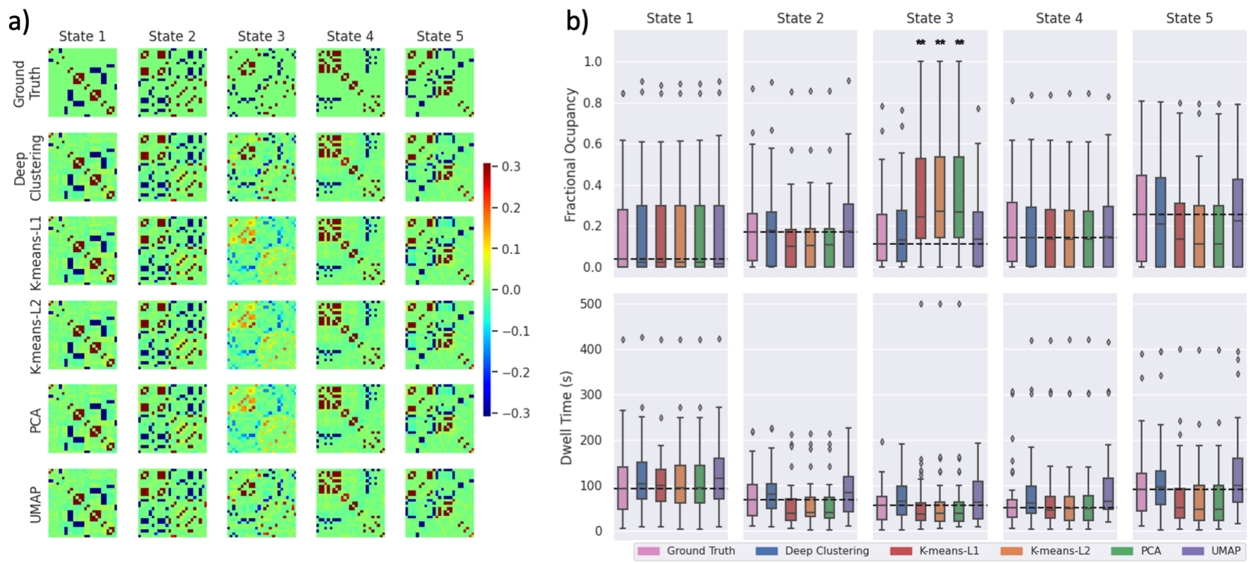


Supplementary Figure 3: Clustering results from one run of each feature selection method applied to synthetic data with low noise. a) State FC matrices are plotted with connectivity indicated by the colour bar. b) The distribution across subjects of fractional occupancy and dwell time measurements are plotted for each state. Boxes show the interquartile range, with a line for the median. The median of the ground truth is shown as a dashed line across each plot for comparison with other methods. Whiskers extend to the range of the data, not including outliers which are shown as diamonds. Significant differences from the true distributions, measured by unpaired two-tailed t-tests, are indicated as follows: *$p < 0.01$, **$p < 0.001$, ***$p < 0.0001$.

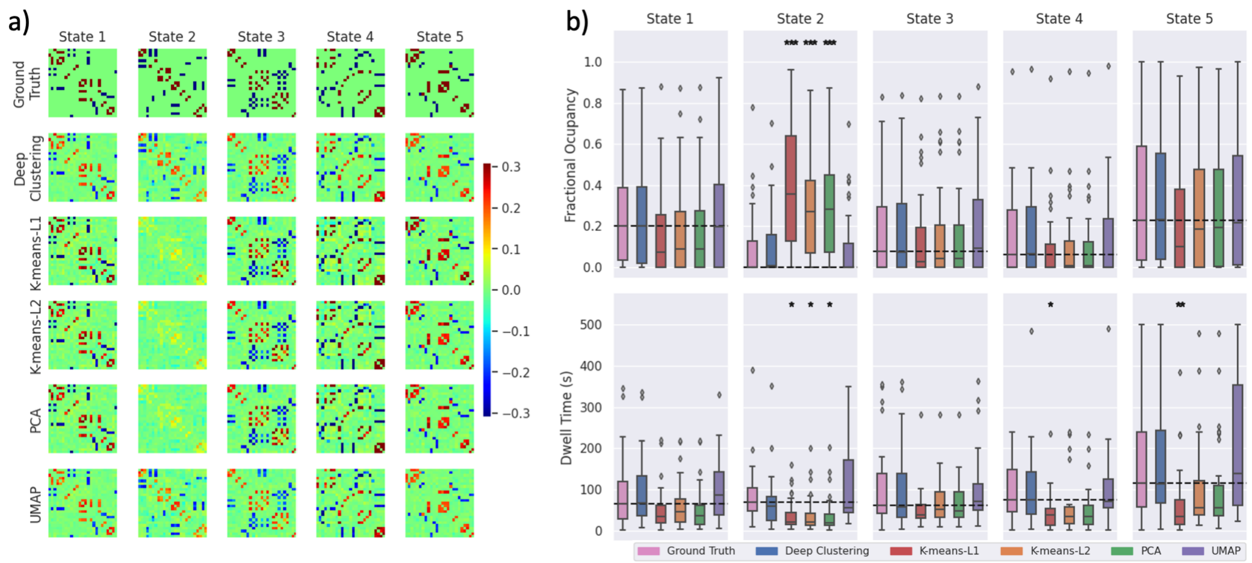


Supplementary Figure 4: Clustering results from one run of each feature selection method applied to synthetic data with medium noise. a) State FC matrices are plotted with connectivity indicated by the colour bar. b) The distribution across subjects of fractional occupancy and dwell time measurements are plotted for each state. Boxes show the interquartile range, with a line for the median. The median of the ground truth is shown as a dashed line across each plot for comparison with other methods. Whiskers extend to the range of the data, not including outliers which are shown as diamonds. Significant differences from the true distributions, measured by unpaired two-tailed t-tests, are indicated as follows: *$p < 0.01$, **$p < 0.001$, ***$p < 0.0001$.

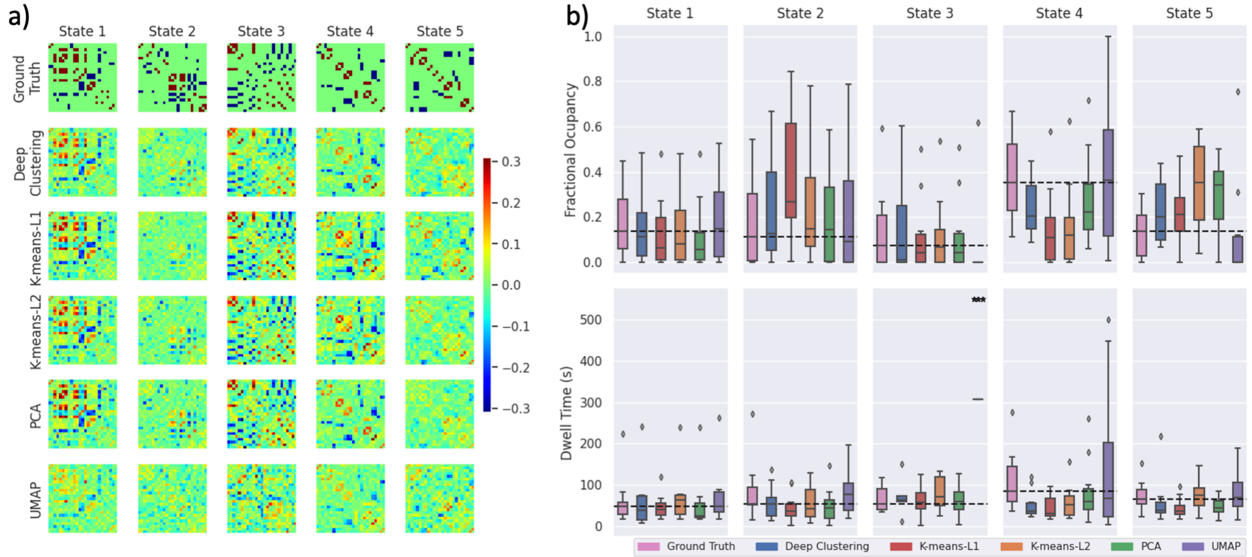


Supplementary Figure 5: Clustering results from one run of each feature selection method applied to synthetic data with 10 subjects. a) State FC matrices are plotted with connectivity indicated by the colour bar. b) The distribution across subjects of fractional occupancy and dwell time measurements are plotted for each state. Boxes show the interquartile range, with a line for the median. The median of the ground truth is shown as a dashed line across each plot for comparison with other methods. Whiskers extend to the range of the data, not including outliers which are shown as diamonds. Significant differences from the true distributions, measured by unpaired two-tailed t-tests, are indicated as follows: $*p < 0.01$, $**p < 0.001$, $***p < 0.0001$.

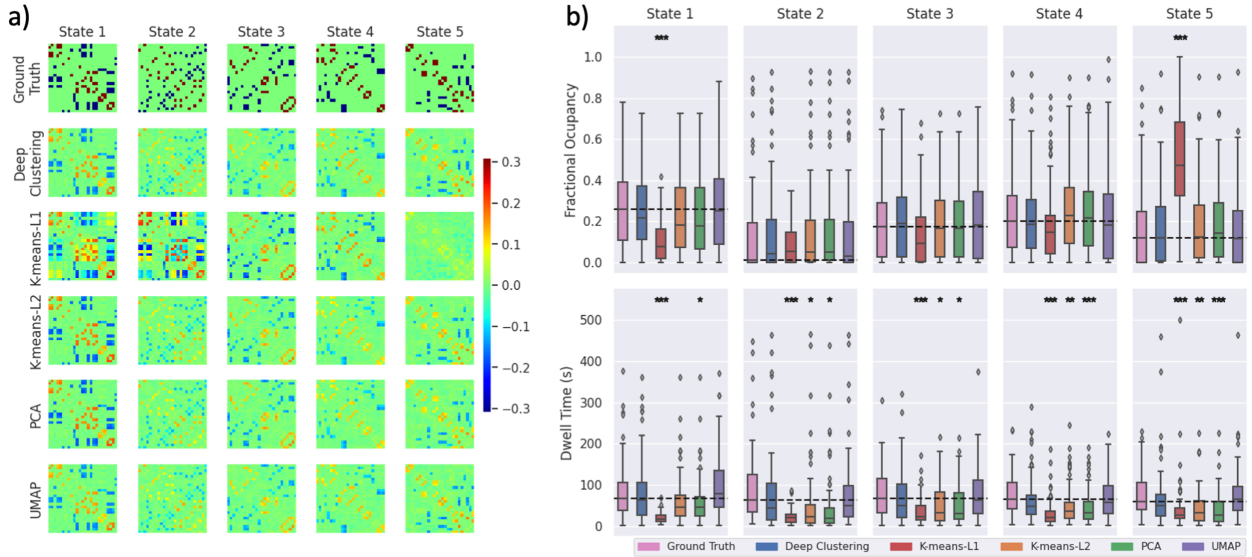


Supplementary Figure 6: Clustering results from one run of each feature selection method applied to synthetic data with 100 subjects. a) State FC matrices are plotted with connectivity indicated by the colour bar. b) The distribution across subjects of fractional occupancy and dwell time measurements are plotted for each state. Boxes show the interquartile range, with a line for the median. The median of the ground truth is shown as a dashed line across each plot for comparison with other methods. Whiskers extend to the range of the data, not including outliers which are shown as diamonds. Significant differences from the true distributions, measured by unpaired two-tailed t-tests, are indicated as follows: $*p < 0.01$, $**p < 0.001$, $***p < 0.0001$.

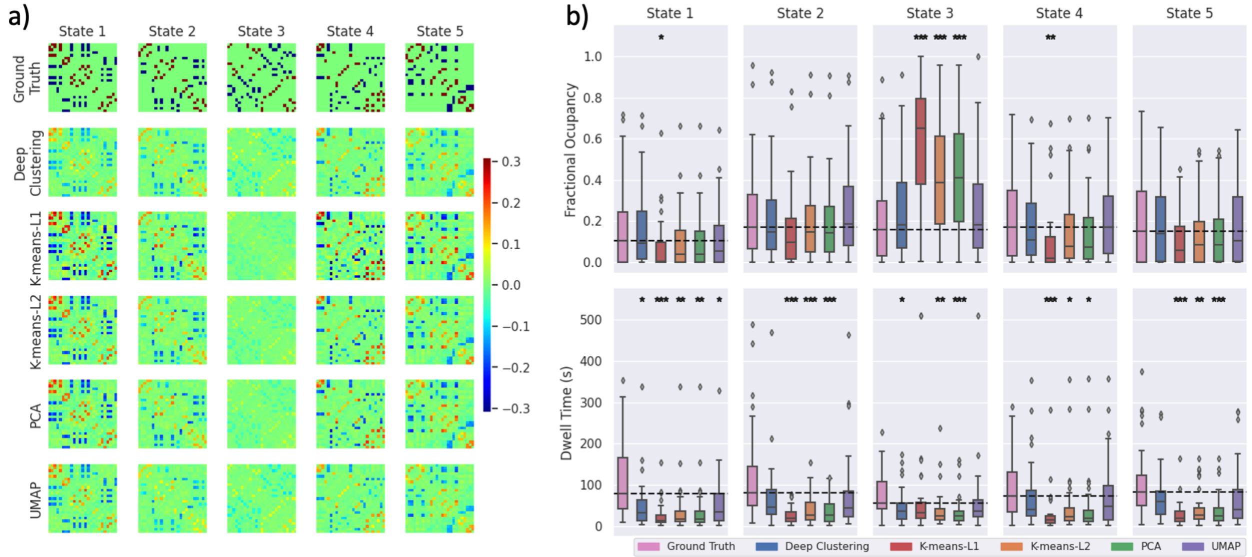


Supplementary Figure 7: Clustering results from one run of each feature selection method applied to synthetic data processed with a 30 s window. a) State FC matrices are plotted with connectivity indicated by the colour bar. b) The distribution across subjects of fractional occupancy and dwell time measurements are plotted for each state. Boxes show the interquartile range, with a line for the median. The median of the ground truth is shown as a dashed line across each plot for comparison with other methods. Whiskers extend to the range of the data, not including outliers which are shown as diamonds. Significant differences from the true distributions, measured by unpaired two-tailed t-tests, are indicated as follows: $^*p < 0.01$, $^{**}p < 0.001$, $^{***}p < 0.0001$.

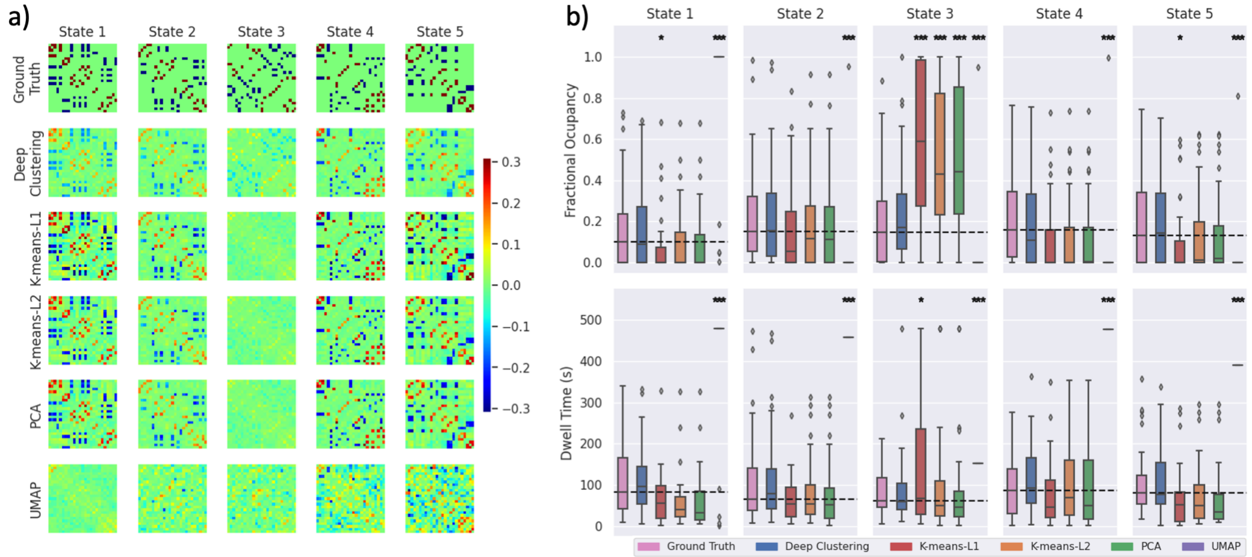


Supplementary Figure 8: Clustering results from one run of each feature selection method applied to synthetic data processed with a 60 s window. a) State FC matrices are plotted with connectivity indicated by the colour bar. b) The distribution across subjects of fractional occupancy and dwell time measurements are plotted for each state. Boxes show the interquartile range, with a line for the median. The median of the ground truth is shown as a dashed line across each plot for comparison with other methods. Whiskers extend to the range of the data, not including outliers which are shown as diamonds. Significant differences from the true distributions, measured by unpaired two-tailed t-tests, are indicated as follows: $^*p < 0.01$, $^{**}p < 0.001$, $^{***}p < 0.0001$.

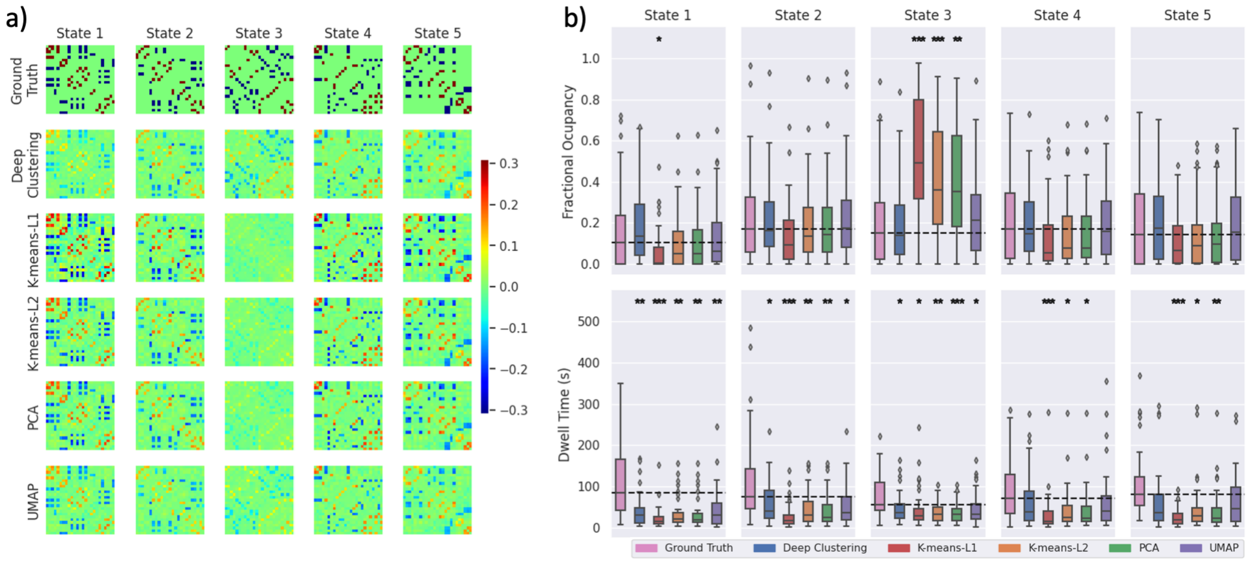


Supplementary Figure 9: Clustering results from one run of each feature selection method applied to synthetic data processed with a Hamming window. a) State FC matrices are plotted with connectivity indicated by the colour bar. b) The distribution across subjects of fractional occupancy and dwell time measurements are plotted for each state. Boxes show the interquartile range, with a line for the median. The median of the ground truth is shown as a dashed line across each plot for comparison with other methods. Whiskers extend to the range of the data, not including outliers which are shown as diamonds. Significant differences from the true distributions, measured by unpaired two-tailed t-tests, are indicated as follows: $*p < 0.01$, $**p < 0.001$, $***p < 0.0001$.

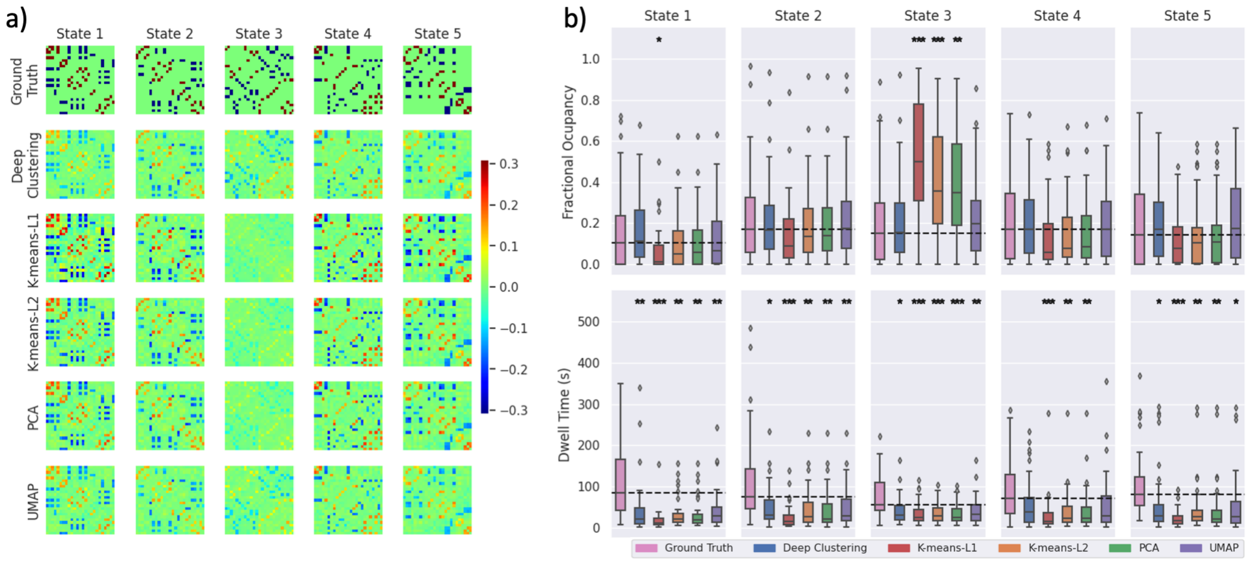


Supplementary Figure 10: Clustering results from one run of each feature selection method applied to synthetic data processed with a Hanning window. a) State FC matrices are plotted with connectivity indicated by the colour bar. b) The distribution across subjects of fractional occupancy and dwell time measurements are plotted for each state. Boxes show the interquartile range, with a line for the median. The median of the ground truth is shown as a dashed line across each plot for comparison with other methods. Whiskers extend to the range of the data, not including outliers which are shown as diamonds. Significant differences from the true distributions, measured by unpaired two-tailed t-tests, are indicated as follows: $*p < 0.01$, $**p < 0.001$, $***p < 0.0001$.

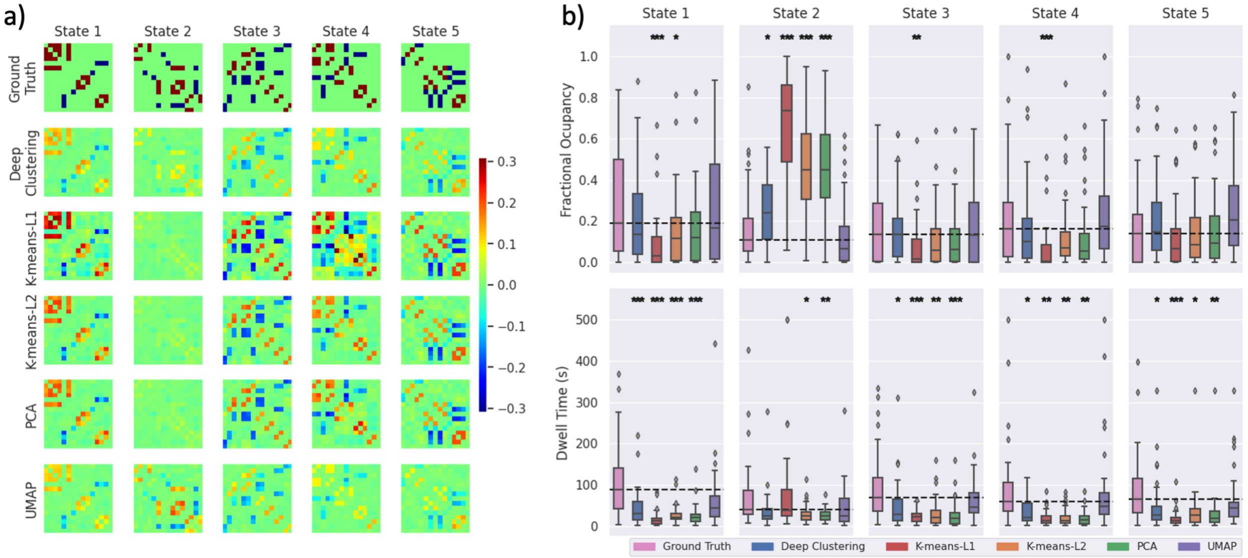


Supplementary Figure 11: Clustering results from one run of each feature selection method applied to synthetic data with 15 nodes. a) State FC matrices are plotted with connectivity indicated by the colour bar. b) The distribution across subjects of fractional occupancy and dwell time measurements are plotted for each state. Boxes show the interquartile range, with a line for the median. The median of the ground truth is shown as a dashed line across each plot for comparison with other methods. Whiskers extend to the range of the data, not including outliers which are shown as diamonds. Significant differences from the true distributions, measured by unpaired two-tailed t-tests, are indicated as follows: $*p < 0.01$, $**p < 0.001$, $***p < 0.0001$.

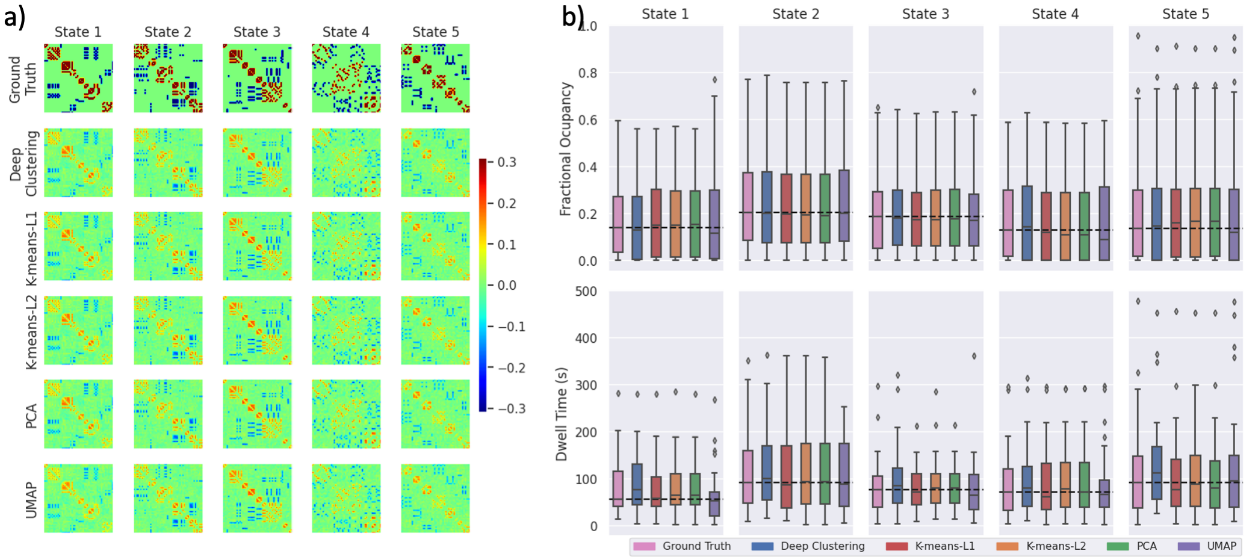


Supplementary Figure 12: Clustering results from one run of each feature selection method applied to synthetic data with 50 nodes. a) State FC matrices are plotted with connectivity indicated by the colour bar. b) The distribution across subjects of fractional occupancy and dwell time measurements are plotted for each state. Boxes show the interquartile range, with a line for the median. The median of the ground truth is shown as a dashed line across each plot for comparison with other methods. Whiskers extend to the range of the data, not including outliers which are shown as diamonds. Significant differences from the true distributions, measured by unpaired two-tailed t-tests, are indicated as follows: $*p < 0.01$, $**p < 0.001$, $***p < 0.0001$.

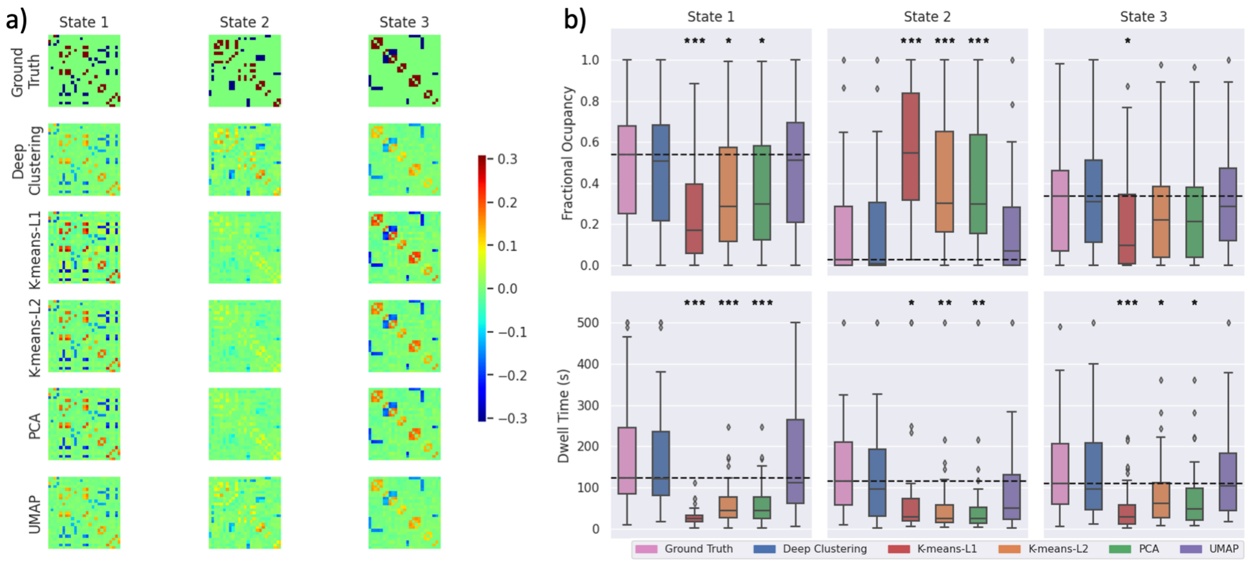


Supplementary Figure 13: Clustering results from one run of each feature selection method applied to synthetic data with 3 states. a) State FC matrices are plotted with connectivity indicated by the colour bar. b) The distribution across subjects of fractional occupancy and dwell time measurements are plotted for each state. Boxes show the interquartile range, with a line for the median. The median of the ground truth is shown as a dashed line across each plot for comparison with other methods. Whiskers extend to the range of the data, not including outliers which are shown as diamonds. Significant differences from the true distributions, measured by unpaired two-tailed t-tests, are indicated as follows: $*p < 0.01$, $**p < 0.001$, $***p < 0.0001$.

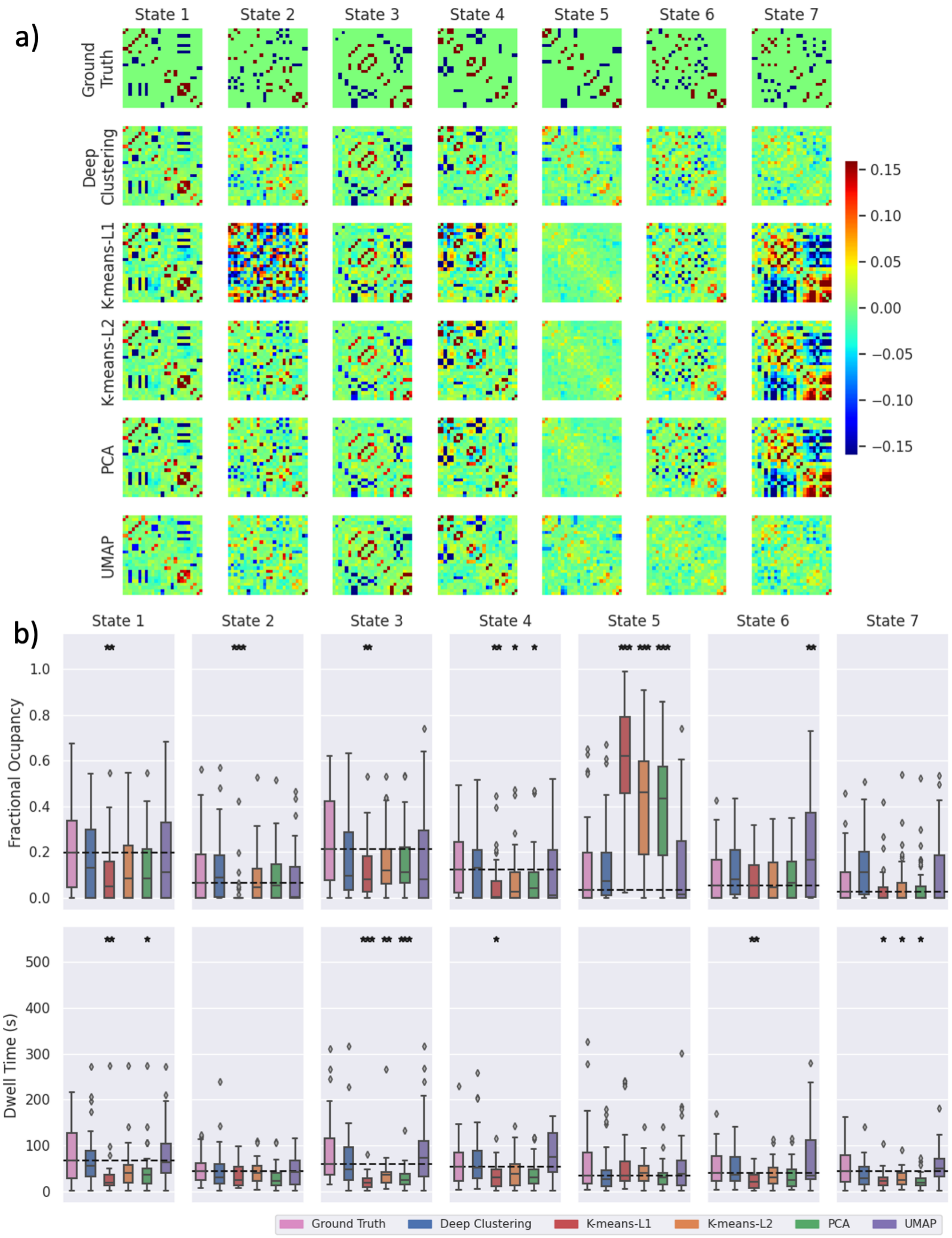


Supplementary Figure 14: Clustering results from one run of each feature selection method applied to synthetic data with 7 states. a) State FC matrices are plotted with connectivity indicated by the colour bar. b) The distribution across subjects of fractional occupancy and dwell time measurements are plotted for each state. Boxes show the interquartile range, with a line for the median. The median of the ground truth is shown as a dashed line across each plot for comparison with other methods. Whiskers extend to the range of the data, not including outliers which are shown as diamonds. Significant differences from the true distributions, measured by unpaired two-tailed t-tests, are indicated as follows: $*p < 0.01$, $**p < 0.001$, $***p < 0.0001$.

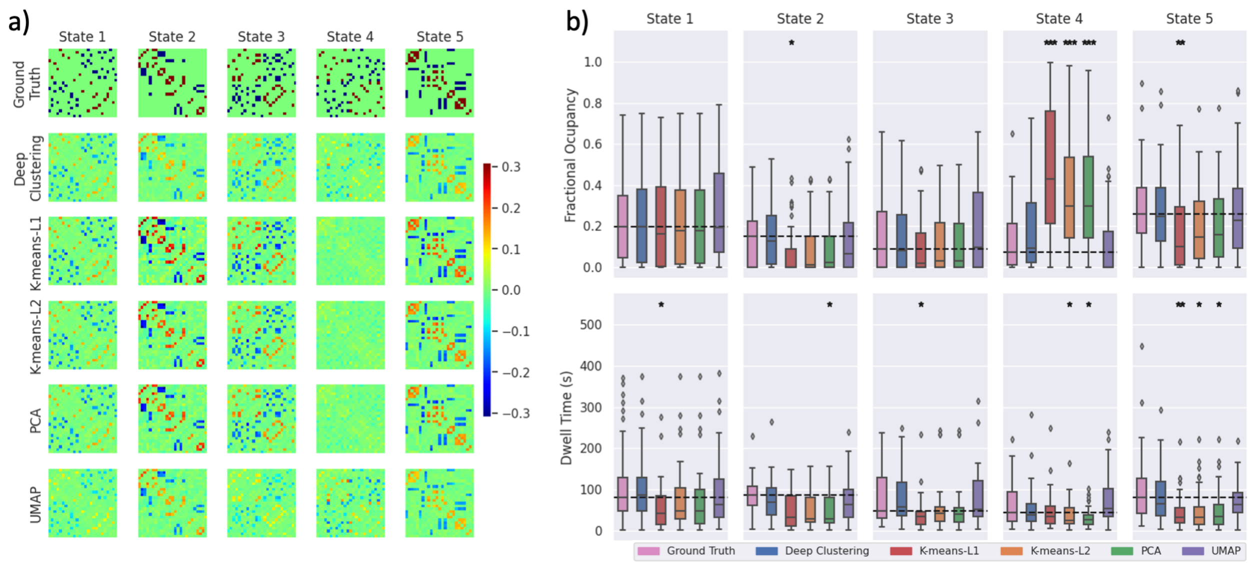


Supplementary Figure 15: Clustering results from one run of each feature selection method applied to synthetic data constructed with the Windkessel-Balloon HRF. a) State FC matrices are plotted with connectivity indicated by the colour bar. b) The distribution across subjects of fractional occupancy and dwell time measurements are plotted for each state. Boxes show the interquartile range, with a line for the median. The median of the ground truth is shown as a dashed line across each plot for comparison with other methods. Whiskers extend to the range of the data, not including outliers which are shown as diamonds. Significant differences from the true distributions, measured by unpaired two-tailed t-tests, are indicated as follows: $*p < 0.01$, $**p < 0.001$, $***p < 0.0001$.

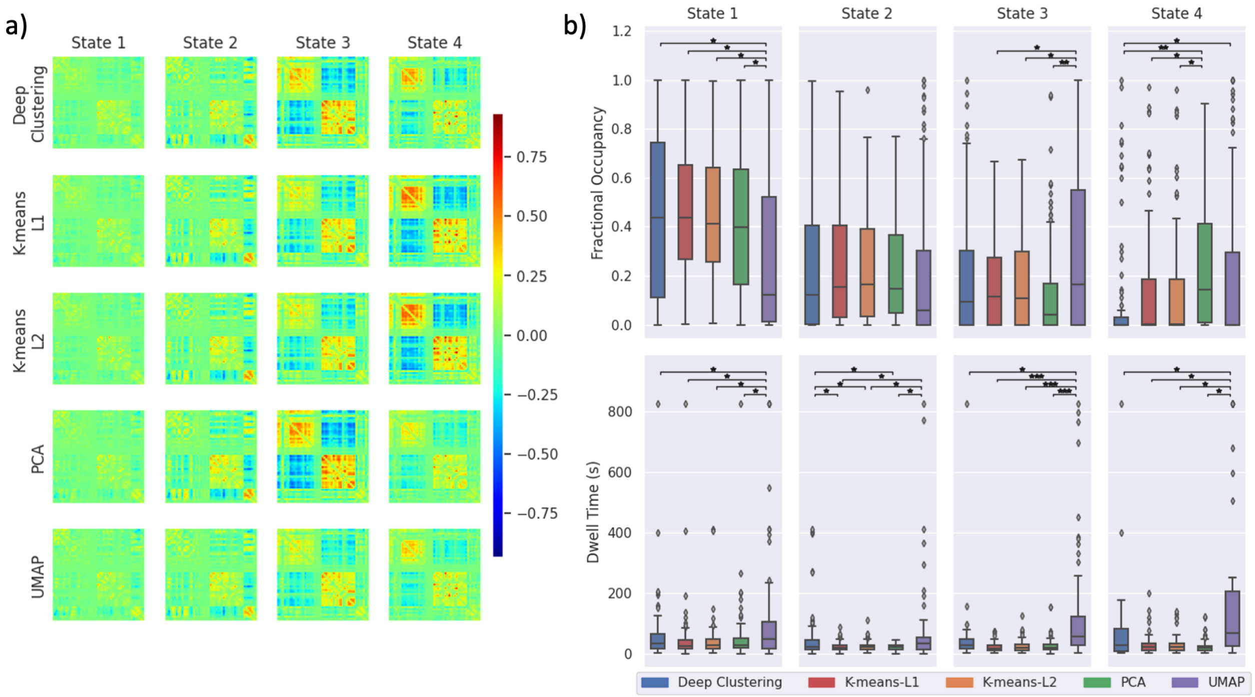


Supplementary Figure 16: Clustering results from each feature selection method applied to the second group of 100 subjects from the Human Connectome Project. a) State FC matrices resulting from each method, with connectivity indicated by the colour bar. b) Fractional occupancy and dwell time measurements across subjects are shown for each state. Boxes show the interquartile range with a line for the median. Whiskers extend to the range of the data, not including outliers which are shown as diamonds. Measurements were compared using a one-way ANOVA, with post hoc pairwise comparison using two-tailed t-tests. FDR-corrected: $*p < 0.05$, $**p < 0.001$, $***p < 0.0001$.

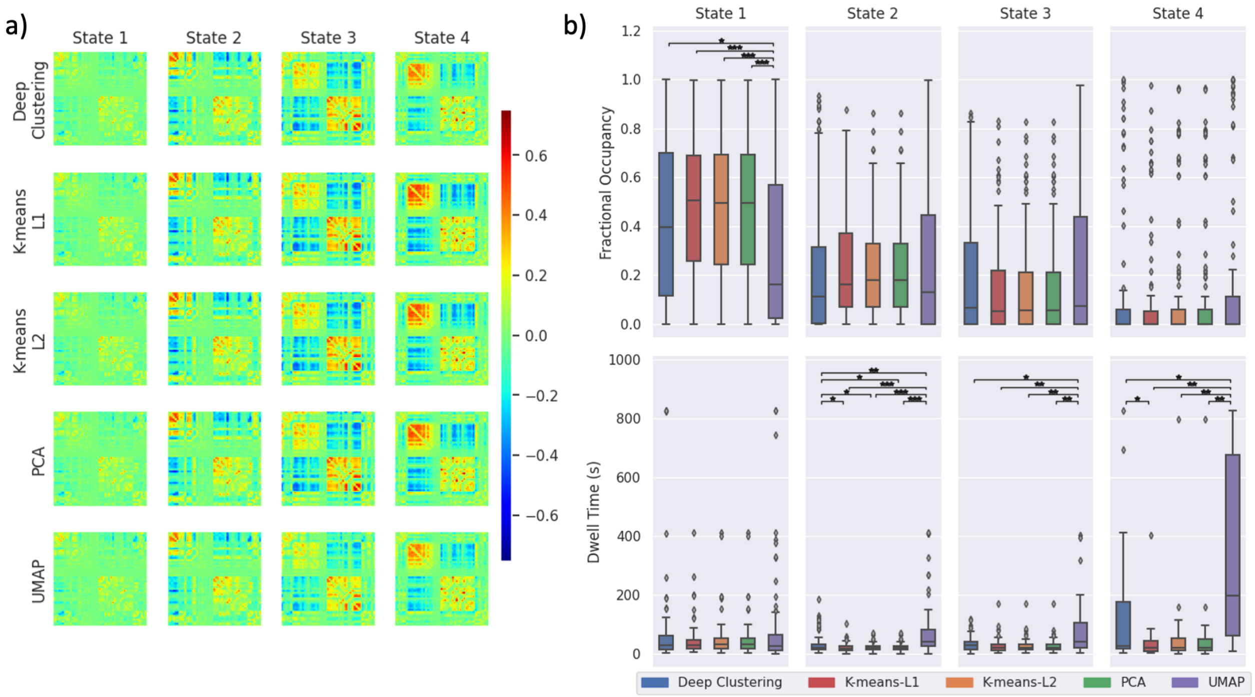


Supplementary Figure 17: Clustering results from each feature selection method applied to the third group of 100 subjects from the Human Connectome Project. a) State FC matrices resulting from each method, with connectivity indicated by the colour bar. b) Fractional occupancy and dwell time measurements across subjects are shown for each state. Boxes show the interquartile range with a line for the median. Whiskers extend to the range of the data, not including outliers which are shown as diamonds. Measurements were compared using a one-way ANOVA, with post hoc pairwise comparison using two-tailed t-tests. FDR-corrected: $*p < 0.05$, $**p < 0.001$, $***p < 0.0001$.

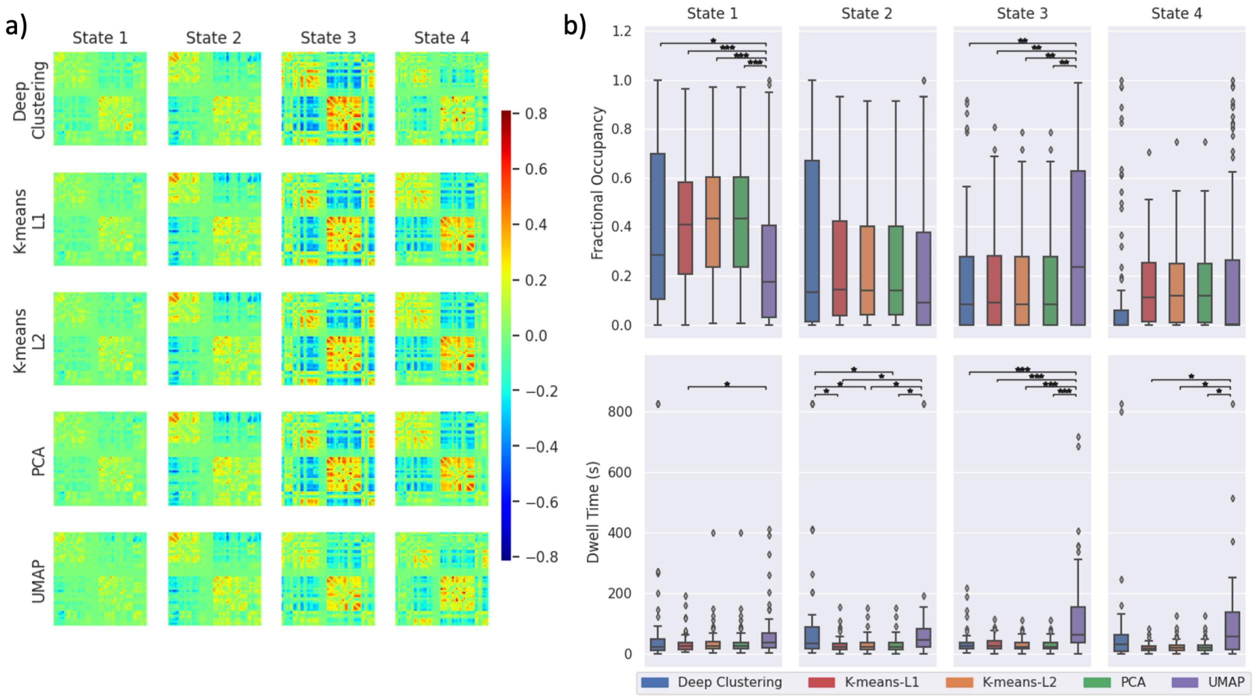


Supplementary Figure 18: Clustering results from each feature selection method applied to the fourth group of 100 subjects from the Human Connectome Project. a) State FC matrices resulting from each method, with connectivity indicated by the colour bar. b) Fractional occupancy and dwell time measurements across subjects are shown for each state. Boxes show the interquartile range with a line for the median. Whiskers extend to the range of the data, not including outliers which are shown as diamonds. Measurements were compared using a one-way ANOVA, with post hoc pairwise comparison using two-tailed t-tests. FDR-corrected: $*p < 0.05$, $**p < 0.001$, $***p < 0.0001$.

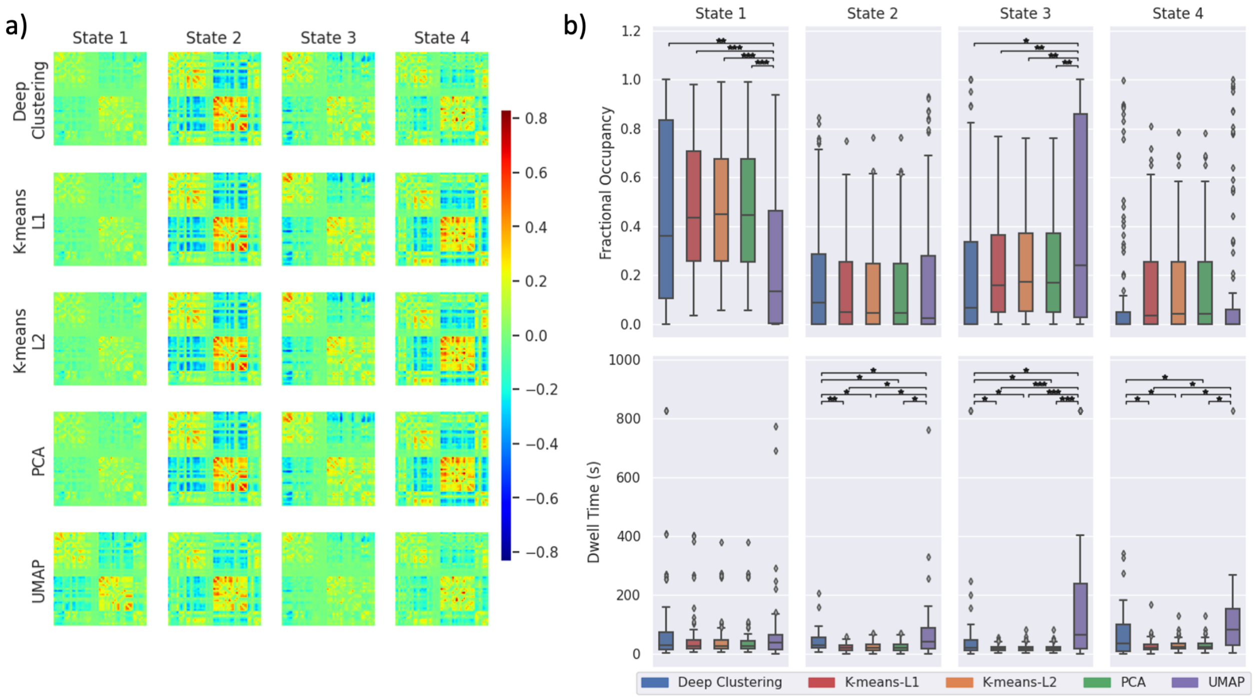


Supplementary Figure 19: Clustering results from each feature selection method applied to the fifth group of 100 subjects from the Human Connectome Project. a) State FC matrices resulting from each method, with connectivity indicated by the colour bar. b) Fractional occupancy and dwell time measurements across subjects are shown for each state. Boxes show the interquartile range with a line for the median. Whiskers extend to the range of the data, not including outliers which are shown as diamonds. Measurements were compared using a one-way ANOVA, with post hoc pairwise comparison using two-tailed t-tests. FDR-corrected: $*p < 0.05$, $**p < 0.001$, $***p < 0.0001$.

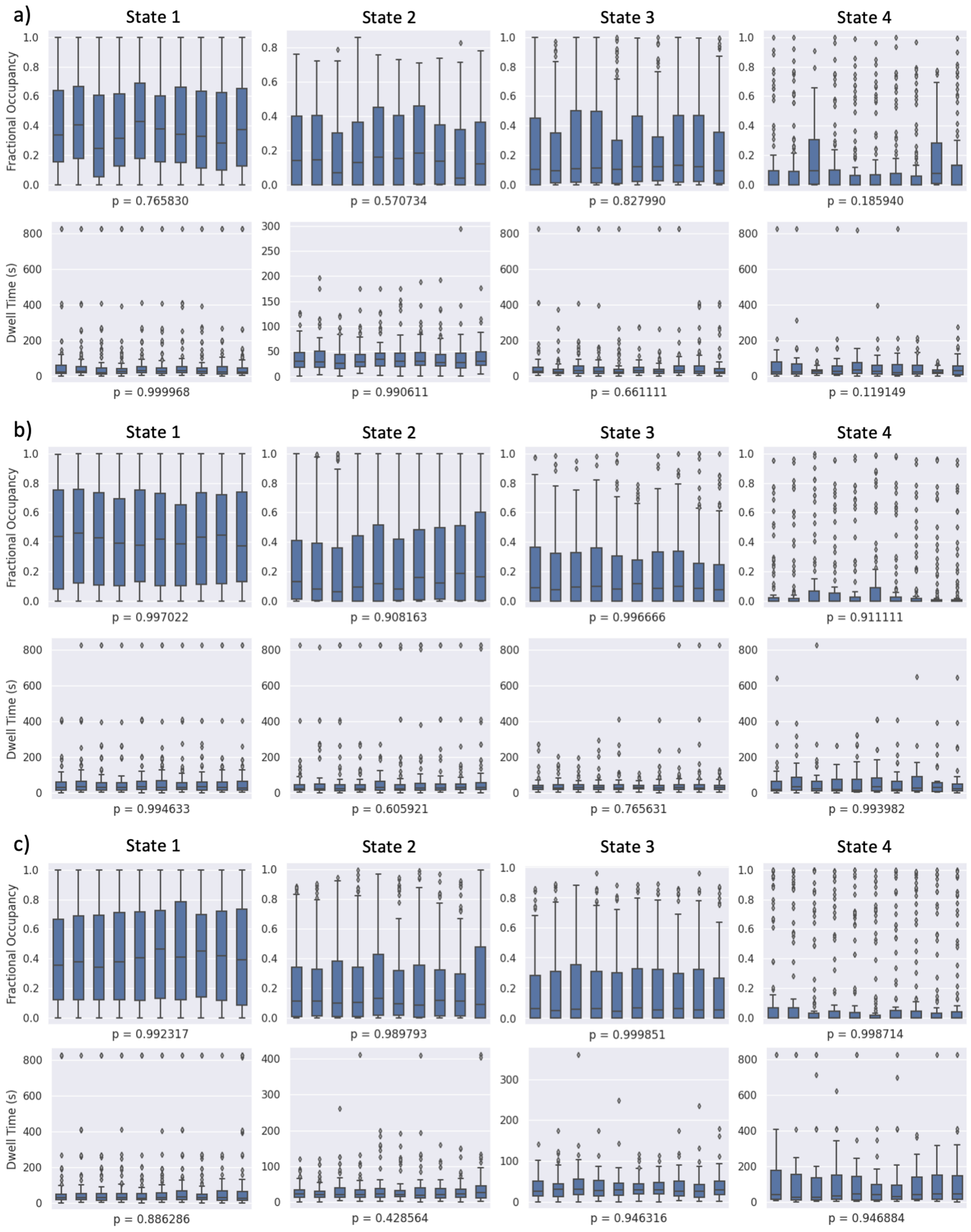


Supplementary Figure 20 (continues on next page)

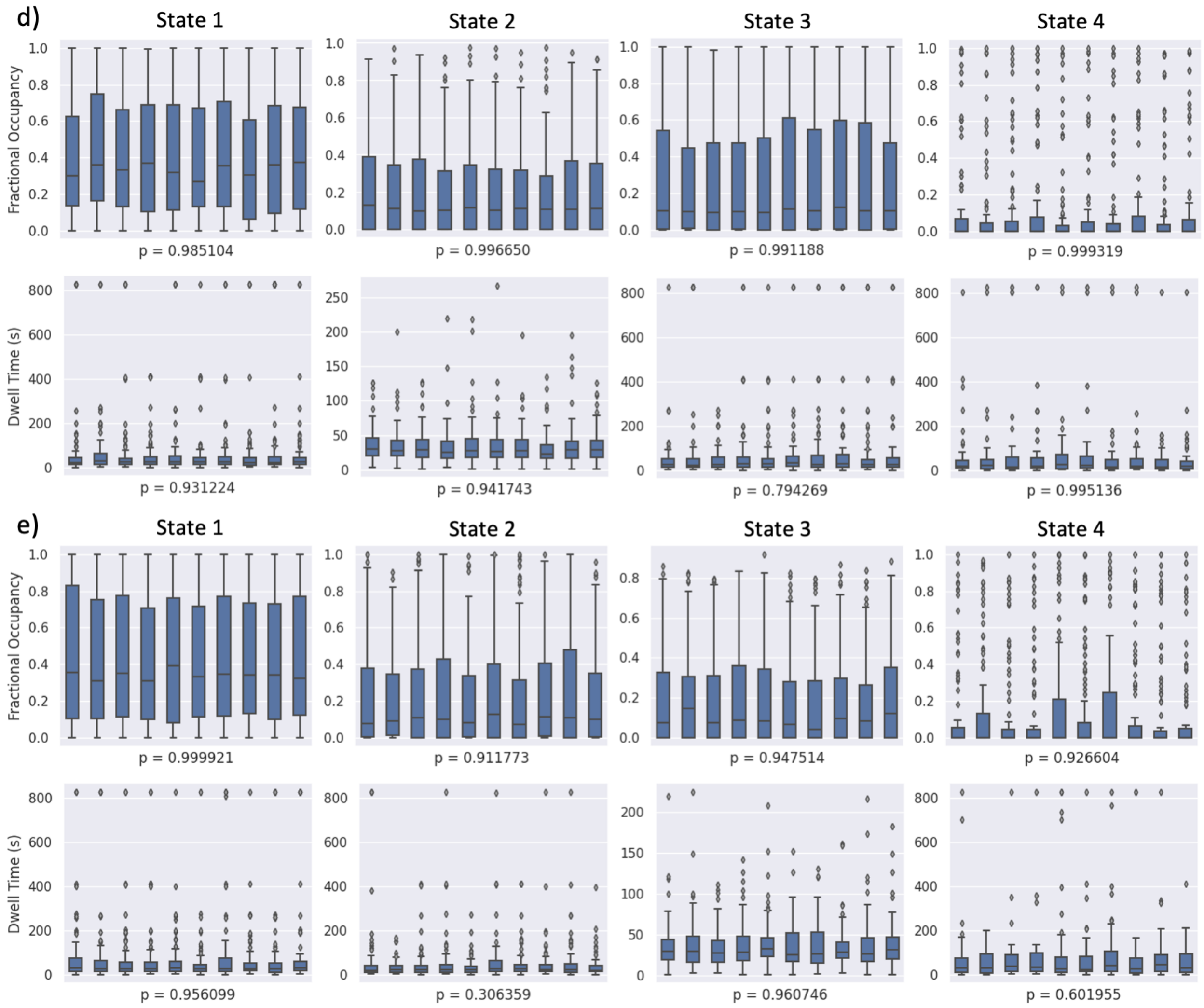


Supplementary Figure 20 (cont.): Repeated runs of deep clustering applied to real data. Each subfigure (a–e) shows results from a different group of 100 participants from the Human Connectome Project. Distributions of fractional occupancy and dwell time are shown for each run of deep clustering (one box per run), with boxes showing the interquartile range with a line for the median and whiskers extending to the range of the data, not including outliers which are shown as diamonds. Each plot shows the p-value of a one-way ANOVA comparing the distribution of measurements across repeated runs. None are significant, indicating that deep clustering gives reproducible measurements of fractional occupancy and dwell time across repeated runs.
